# Comparative phylogeography reveals widespread cryptic diversity driven by ecology in Panamanian birds

**DOI:** 10.1101/2023.03.15.530646

**Authors:** J. F. McLaughlin, Celestino Aguilar, Justin M. Bernstein, Wayra G. Navia-Gine, Luis E. Cueto-Aparicio, Ashleigh C. Alarcon, Brandon D. Alarcon, Rugger Collier, Anshule Takyar, Sidney J. Vong, Oscar G. López-Chong, Robert Driver, Jose R. Loaiza, Luis F. De León, Kristin Saltonstall, Sara E. Lipshutz, Dahiana Arcila, Kinsey M. Brock, Matthew J. Miller

## Abstract

Widespread species often harbor unrecognized genetic diversity, and investigating the factors associated with such cryptic variation can help us better understand the forces driving diversification. Here, we identify potential cryptic species based on a comprehensive dataset of COI mitochondrial DNA barcodes from 2,333 individual Panamanian birds across 429 species, representing 391 (59%) of the 659 resident landbird species of the country, as well as opportunistically sampled waterbirds. We complement this dataset with additional publicly available mitochondrial loci, such as ND2 and cytochrome *b,* obtained from whole mitochondrial genomes from 20 taxa. Using barcode identification numbers (BINs), we find putative cryptic species in 19% of landbird species, highlighting hidden diversity in the relatively well-described avifauna of Panama. Whereas some of these mitochondrial divergence events corresponded with recognized geographic features that likely isolated populations, such as the Cordillera Central highlands, the majority (74%) of lowland splits were between eastern and western populations. The timing of these splits are not temporally coincident across taxa, suggesting that historical events, such as the formation of the Isthmus of Panama and Pleistocene climatic cycles, were not the primary drivers of cryptic diversification. Rather, we observed that forest species, understory species, insectivores, and strongly territorial species—all traits associated with lower dispersal ability—were all more likely to have multiple BINs in Panama, suggesting strong ecological associations with cryptic divergence. Additionally, hand-wing index, a proxy for dispersal capability, was significantly lower in species with multiple BINs, indicating that dispersal ability plays an important role in generating diversity in Neotropical birds. Together, these results underscore the need for evolutionary studies of tropical bird communities to consider ecological factors along with geographic explanations, and that even in areas with well-known avifauna, avian diversity may be substantially underestimated.

**LAY SUMMARY:**

- What factors are common among bird species with cryptic diversity in Panama? What role do geography, ecology, phylogeographic history, and other factors play in generating bird diversity?
- 19% of widely-sampled bird species form two or more distinct DNA barcode clades, suggesting widespread unrecognized diversity.
- Traits associated with reduced dispersal ability, such as use of forest understory, high territoriality, low hand-wing index, and insectivory, were more common in taxa with cryptic diversity.

**Filogeografía comparada revela amplia diversidad críptica causada por la ecología en las aves de Panamá**

**RESUMEN:** Especies extendidas frecuentemente tiene diversidad genética no reconocida, y investigando los factores asociados con esta variación críptica puede ayudarnos a entender las fuerzas que impulsan la diversificación. Aquí, identificamos especies crípticas potenciales basadas en un conjunto de datos de códigos de barras de ADN mitocondrial de 2,333 individuos de aves de Panama en 429 especies, representando 391 (59%) de las 659 especies de aves terrestres residentes del país, además de algunas aves acuáticas muestreada de manera oportunista. Adicionalmente, complementamos estos datos con secuencias mitocondriales disponibles públicamente de otros loci, tal como ND2 o citocroma b, obtenidos de los genomas mitocondriales completos de 20 taxones. Utilizando los números de identificación de código de barras (en ingles: BINs), un sistema taxonómico numérico que proporcina una estimación imparcial de la diversidad potencial a nivel de especie, encontramos especies crípticas putativas en 19% de las especies de aves terrestres, lo que destaca la diversidad oculta en la avifauna bien descrita de Panamá. Aunque algunos de estos eventos de divergencia conciden con características geográficas que probablemente aislaron las poblaciones, la mayoría (74%) de la divergencia en las tierras bajas se encuentra entre las poblaciones orientales y occidentales. El tiempo de esta divergencia no coincidió entre los taxones, sugiriendo que eventos históricos tales como la formación del Istmo de Panamá y los ciclos climáticos del pleistoceno, no fueron los principales impulsores de la especiación. En cambio, observamos asociaciones fuertes entre las características ecológicas y la divergencia mitocondriale: las especies del bosque, sotobosque, con una dieta insectívora, y con territorialidad fuerte mostraton múltiple BINs probables. Adicionalmente, el índice mano-ala, que está asociado a la capacidad de dispersión, fue significativamente menor en las especies con BINs multiples, sugiriendo que la capacidad de dispersión tiene un rol importamente en la generación de la diversidad de las aves neotropicales. Estos resultos demonstran la necesidad de que estudios evolutivos de las comunidades de aves tropicales consideren los factores ecológicos en conjunto con las explicaciones geográficos.

*Palabras clave:* biodiversidad tropical, biogeografía, códigos de barras, dispersión, especies crípticas

## INTRODUCTION

Identifying the number of species in a given region is key to understanding the evolution and maintenance of biodiversity (Bickford *et al*. 2007; Allendorf *et al*. 2009; Pérez-Ponce de León and Nadler 2010). Yet characterizing species diversity remains a crucial challenge in biodiversity research (De León *et al*. 2023), and cryptic species often go unrecognized as distinct due to their high similarity (Fouquet *et al*. 2007; Yan *et al*. 2018; Chenuil *et al*. 2019; Levy and Cox 2020). Despite this difficulty, the identification of cryptic species may illuminate evolutionary processes that generate biodiversity (Campbell, Braile, and Winker 2016; Bickford *et al*. 2007; Barreira, Lijtmaer, and Tubaro 2016; Struck *et al*. 2018). The investigation of cryptic species is often framed as an issue of not adequately describing and cataloging variation (de León and Nadler 2010; Korshunova *et al*. 2017), but studies of cryptic taxa present an opportunity to understand why some species diversify while remaining morphologically conserved (Roux *et al*. 2016; Pulido-Santacruz, Aleixo, and Weir 2018). By linking the diversity of cryptic variation with geography, history, ecology, or a combination of those factors, researchers can better understand what drives the generation of underestimated variation in cryptic species (Bickford *et al*. 2007; Struck *et al*. 2018). Considered together, these sources of variation can provide valuable insights into speciation and diversification, with implications for conservation decision-making (Bickford *et al*. 2007; Saitoh *et al*. 2015; Campbell, Braile, and Winker 2016; Struck *et al*. 2018).

Cryptic diversity is of particular interest in regions with high levels of biodiversity. The Neotropics are home to incredible avian species diversity, with around one in four global bird species found in the region (Haffer 1985; Orme *et al*. 2005). However, this is likely an underestimate, given that studies frequently find species-level genetic diversity, even within recognized species (Tavares *et al*. 2011; Milá *et al*. 2012; Rheindt, Cuervo, and Brumfield 2013; Mendoza *et al*. 2016). This is often the case in widely distributed species, which can harbor extensive genetic variation characteristic of species complexes, such as in *Lepidothrix* (Cheviron, Hackett, and Capparella 2005), *Manacus* (Brumfield *et al*. 2008), *Habia* (Lavinia *et al*. 2015;

Ramírez-Barrera *et al*. 2018, 2019), *Pachyramphus* (Musher and Cracraft 2018), *Malacoptila* (Ferreira *et al*. 2017)*, Arremon* (Cadena, Klicka, and Ricklefs 2007; Navarro-Sigüenza *et al*. 2008; Cadena and Cuervo 2010) and *Phaeothlypis* (Lovette 2004). This underestimation of avian biodiversity across hampers our understanding of the processes that have made the Neotropics such an important hotspot of species diversification (Bickford *et al*. 2007), and it has practical implications for conservation (Bickford *et al*. 2007; Allendorf *et al*. 2009; Valentini, Pompanon, and Taberlet 2009; Lohman *et al*. 2010; Funk, Caminer, and Ron 2012; Crawford *et al*. 2013; Gonçalves *et al*. 2015; Mendoza *et al*. 2016). Although there is some risk that the search for cryptic species may lead to oversplitting and taxonomic inflation (Chaitra, Vasudevan, and Shanker 2004; Isaac, Mallet, and Mace 2004; Hundsdoerfer *et al*. 2019; Chan *et al*. 2020), evidence in birds suggests that their diversity is underestimated (Sangster 2009; Barrowclough *et al*. 2016). Moreover, even if such cryptic taxa may not represent distinct species, they are distinct populations on their own evolutionary trajectories, and identifying such population-level differences can allow us to understand different parts of the speciation continuum.

Mitochondrial barcoding provides us with a powerful tool to detect cryptic diversity (Arnot, Roper, and Bayoumi 1993; Floyd *et al*. 2002; Hebert, Ratnasingham, and deWaard 2003; Hebert and Gregory 2005; Clare *et al*. 2007; Vasconcelos *et al*. 2016; Imtiaz, Nor, and Naim 2017; Bernstein *et al*. 2021). Previous efforts using mitochondrial markers have documented cryptic variation in multiple Panamanian birds (González *et al*. 2003; Miller *et al*. 2008; Miller *et al*. 2011; Bryson *et al*. 2014; Loaiza *et al*. 2016; Lopez *et al*. 2016). However, as useful as single-taxon studies are, they provide only one example of potential widespread patterns of phylogeographic diversity. Yet, mitochondrial barcoding, with its relative ease of locus acquisition and low cost, provides a simple but powerful tool to build large scale comparative datasets across a wide range of taxa (Kerr *et al*. 2007, 2009; Pereira *et al*. 2013; Mendoza *et al*. 2016). Mitochondrial barcoding is particularly helpful due to the same locus having been sequenced in a wide range of taxa (Bronstein, Kroh, and Haring 2018). With comparative datasets, we can better estimate the occurrence of cryptic species, which in turn allows us to better understand the origin of biodiversity in a given region.

Barcoding many species exhibiting cryptic variation can detect patterns in where species turnovers occur geographically (e.g., such as corresponding with known suture zones in other taxa) or what ecological factors are more common in cryptically diverse taxa (Bickford *et al*. 2007; Struck *et al*. 2018). Addressing the vast biodiversity of the Neotropics necessitates large comparative phylogeographic datasets (Kerr *et al*. 2009; Tavares *et al*. 2011; Miller *et al*. 2021). Hypotheses on the origin of Neotropical diversity tend to fall into a few broad groups. First, biogeographic explanations explain how the geologic and environmental history of a region has created isolated populations that lead to diversification (Sick 1967; Haffer 1969, 1985, 1997; Bush 1994; Sedano and Burns 2010; Smith *et al*. 2014; Ferreira *et al*. 2017). In Panama, these are further grouped into those emphasizing the process of the formation of the Isthmus of Panama (DaCosta and Klicka 2008; Smith and Klicka 2010; Leigh, O’Dea, and Vermeij 2014a), and those emphasizing paleoclimatic fluctuations driving a shifting mosaic of forest and savannah across the landscape (Smith, Amei, and Klicka 2012). Secondly, there are ecological explanations which focus on the role of competition and diversification, with the profusion of niches driving the diversification of species to fill them (Klopfer and MacArthur 1960, 1961; Emerson and Kolm 2005; Brown 2014; Moles and Ollerton 2016). While both explanations likely play roles in generating Panamanian biodiversity (Bush 1994; Smith *et al*. 2014), the question of which prevails in a given region and at finer spatial and temporal scales remains unclear.

By DNA barcoding multiple taxa across wide geographic ranges, we can test hypotheses of both biogeographic and ecological divergence. The Panamanian region is a topographically and ecologically diverse region, especially considering its small size (Ridgely and Gwynne 1992; Siegel and Olson 2008; Angehr and Dean 2010). We expect that Panama’s many islands and disjunct highlands (Figure 1A) will harbor a disproportionate number of cryptic species, as has been found elsewhere (Saitoh *et al*. 2015; Campbell, Braile, and Winker 2016). Both the communities of these discrete regions and the more continuously distributed lowland taxa may have been subject to historic isolation, especially prior to the final closure of the Isthmus of Panama approximately 2.7 to 4.2 million years ago (Leigh, O’Dea, and Vermeij 2014b; O’Dea *et al*. 2016; Jaramillo *et al*. 2017), or by the possible expansion of savannah habitats and formation of forest refugia during the Pleistocene (Smith, Amei, and Klicka 2012). In particular, lowland Panama has been recognized as a hotspot for species turnover—replacement of a given taxa with other taxa— in birds (Miller, Bermingham, and Ricklefs 2007; Miller *et al*. 2011; Loaiza *et al*. 2016; Lopez *et al*. 2016; McLaughlin, Garzón, *et al*. 2020), as well as in freshwater fish (Bermingham and Martin 1998; Martin and Bermingham 2000; Perdices *et al*. 2002; Smith and Bermingham 2005; Bagley and Johnson 2014), mammals (Cortés-Ortiz *et al*. 2003), herpetofauna (Crawford 2003; Crawford and Smith 2005; Bagley and Johnson 2014), insects (Bagley and Johnson 2014; Eskildsen *et al*. 2018), and plants (Dick, Abdul-Salim, and Bermingham 2003). Thus, using barcoding, we can investigate if biogeography is an important factor in generating avian diversity, by testing if divergence events are broadly coincident in time (Naka and Brumfield 2018). Beyond biogeographic explanations, our diverse sampling (Figure 2) allows us to investigate whether specific ecological traits, such as habitat preference (De León *et al*. 2010; Zhang *et al*. 2012; Harvey *et al*. 2017), dispersal ability (Claramunt *et al*. 2012; Weeks and Claramunt 2014; Crouch *et al*. 2019), territoriality (Tobias *et al*. 2016), and diet (Sheard *et al*. 2020; Miller *et al*. 2021) are associated with cryptic divergence.

**Figure 1:**
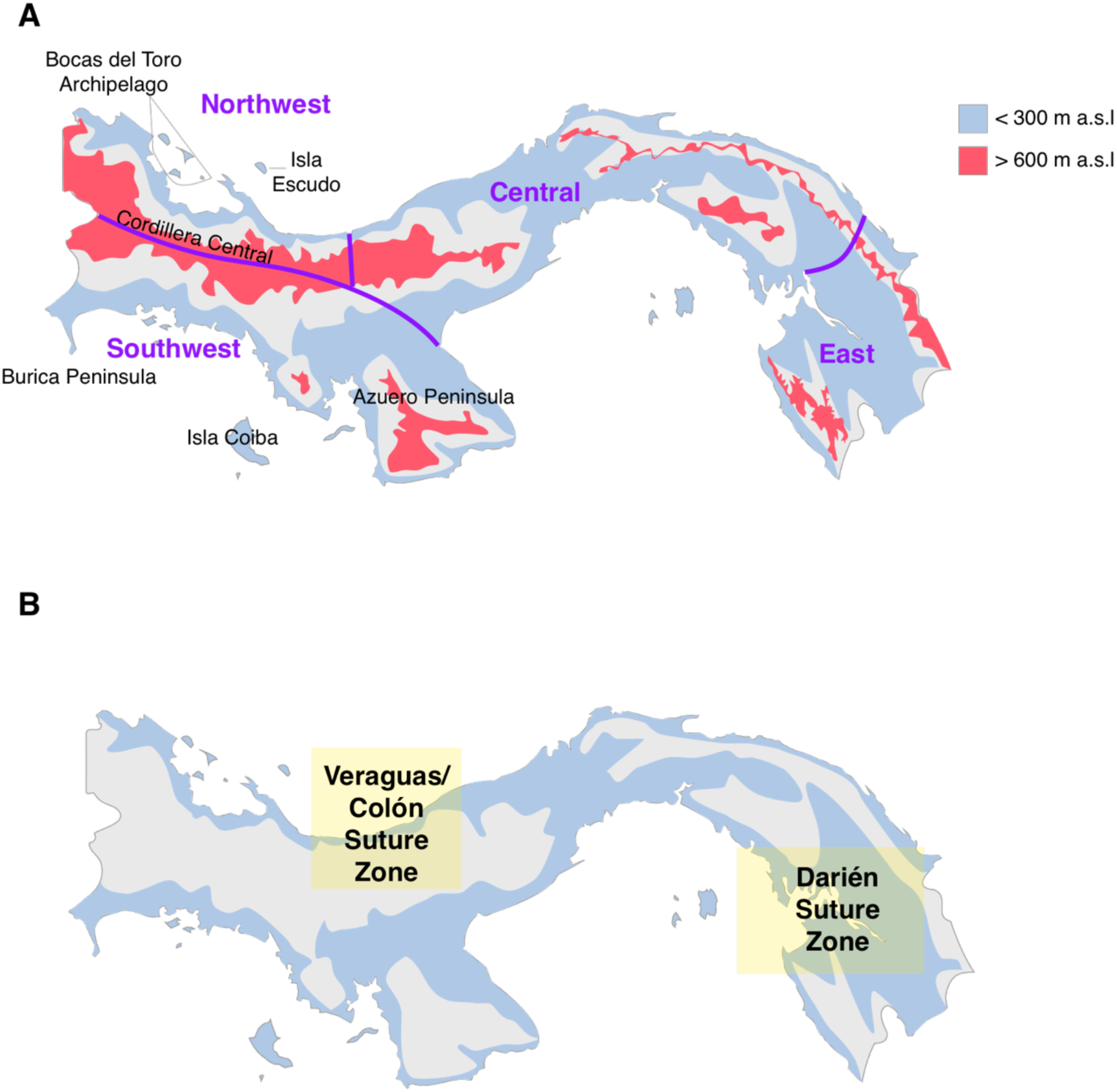
Map of Panama indicating major geographic features considered in this study. A) Pink areas are highland regions with greater than 600 meters above sea levels (m.a.s.l) elevation, while light blue show those with less than 300 m.a.s.l. Purple lines indicate the broad geographic areas used to define sampling regions, with names in purple. B) Lowland suture zones. Light yellow shading indicates general locations of suture zones described in this study.

**Figure 2:**
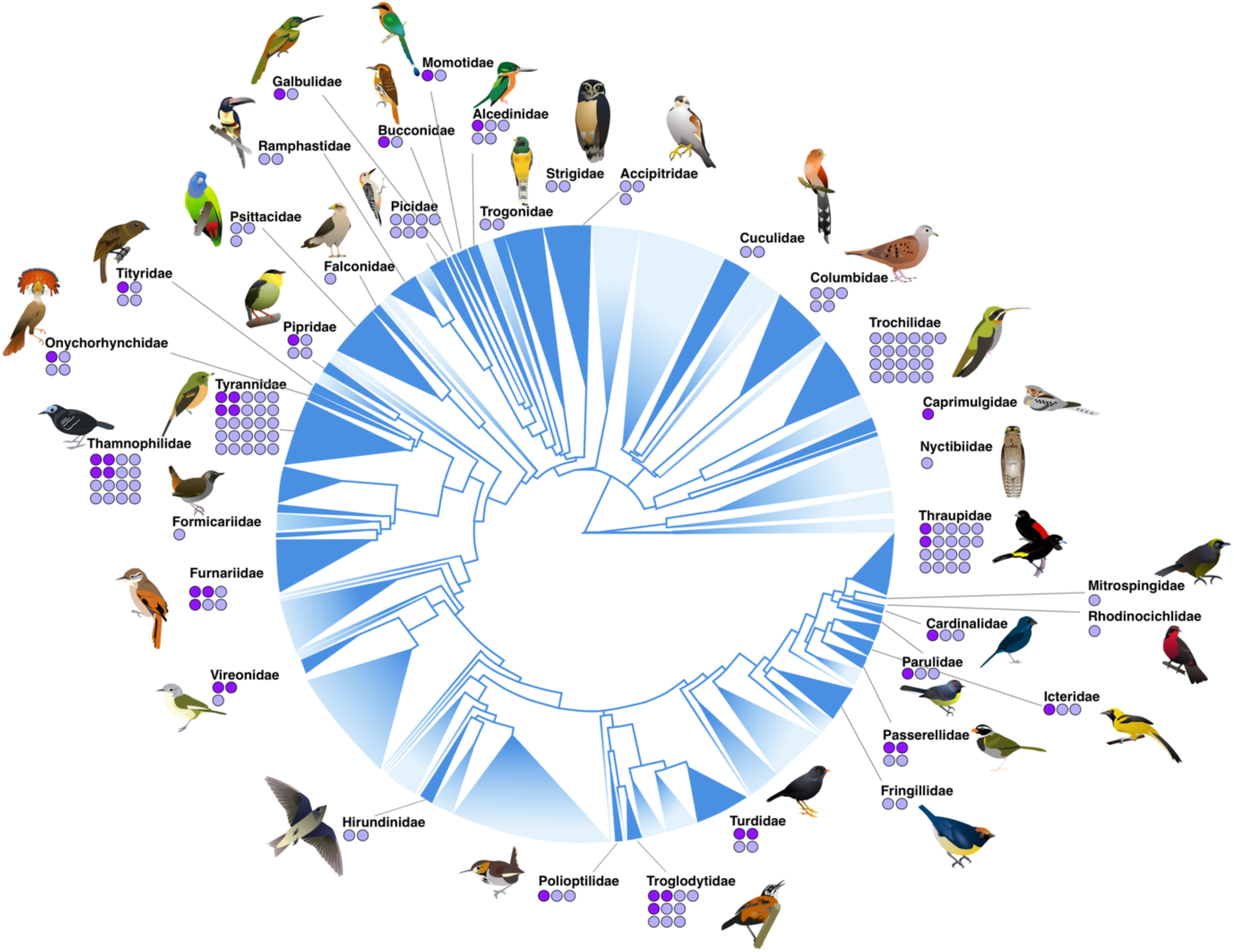
Phylogenetic sampling of Panamanian birds. Overall avian tree of life modified from Jetz *et al* (2012). Sampled families are shown in dark blue, with unsampled lineages shown with topology faded out; size of clade scaled by number of species. For each sampled family, dots indicate the number of taxa sampled across multiple regions of Panama (one dot per sampled species). Light purple dots indicate lineages with a single mitochondrial BIN; dark purple indicate those with two or more.

We set out to investigate the connections between geography, ecology, and the occurrence of cryptic phylogeographic variation indicative of species with a multifaceted mitochondrial dataset. We barcoded 429 species of birds, including 391 landbird species (59%) of the 659 documented resident landbirds in the country. We found that specific ecological traits, including dispersal ability, territoriality, diet, and habitat, were significantly over-represented in taxa with mitochondrial breaks, suggesting these may strongly contribute to the diversification of Neotropical birds.

## METHODS

### COI Barcode Survey of Panamanian Birds

We developed a COI barcode dataset of 2,333 birds from across Panama as part of sequencing for the Barcode of Life Database (BOLD; Ratnasingham and Hebert 2007, 2013). Our dataset includes 429 species as defined in the original taxon dataset from BOLD; however, Angehr and Dean 2010 define 484 by splitting several taxa that are not in the taxonomy used in BOLD. The majority of these birds were sequenced through the Smithsonian Institution’s (SI) Barcode of Life initiative (Schindel *et al*. 2011) and the Smithsonian Tropical Research Institute (STRI; original data presented here). With a few exceptions, every sequence in the SI and STRI barcoding datasets is represented by a museum voucher specimen (Table A1). We determined whether a species had mitochondrial splits—i.e., had 2 or more barcode clusters defined in BOLD—by using the barcode index number (BIN; Ratnasingham and Hebert 2007, 2013) as implemented in BOLD via alignment and clustering of COI sequences. In this method, individuals that are more than twice the distance of divergence within a cluster being taken as the start of a new cluster, followed by use of a Markovian analysis to refine clusters (Ratnasingham and Hebert 2013). Key benefits of this method are in the ease and low-cost of the method (Tavares *et al*. 2011; Milá *et al*. 2012), and the relatively high reliability in assigning individuals to species in past studies (Yoo *et al*. 2006; Kerr *et al*. 2007). It does carry the standard limitations of any single-marker method of evaluating diversity, namely that a single locus may not be reflective of the total evolutionary history of a taxon. However, mitochondrial studies are still valuable where large-scale nuclear sequencing is not feasible, and are useful in determining where to focus with more in-depth sequencing efforts.

### Improving Geographic Coverage Through Mitogenomic Haplotyping

Though covering over 2000 birds, our COI database does not fully capture available data on the distribution of mitochondrial diversity and structure in Panamanian birds that is available either as part of previously published studies, (e.g. Miller *et al*. 2010; Smith *et al*. 2014; Miller *et al*. 2021). Because mitochondrial DNA is non-recombining, whole mitochondrial genomes can connect disparate mtDNA datasets into congruent haplogroups, functioning as a “Rosetta Stone” to leverage multiple mitochondrial loci into a large common dataset. As part of several long-term projects on the comparative genomics of Panamanian lowlands, we filtered mtDNA reads from whole genome shotgun sequencing (e.g. do Amaral *et al*. 2015) for 20 taxa identified with distinct COI BINs in Panama of resident lowland birds sampled in western (Bocas del Toro) and eastern (Darién) Panama (Table 1).We sequenced two individuals from each of those populations, preparing genomic libraries with the NEB Ultra II protocol and sequencing them on an Illumina NovaSeq. We then used bbduk, a utility within the bbmap program (Bushnell 2014), to trim and perform initial quality control on reads. We then downloaded 215 additional mitochondrial sequences, including ND2, cytB, ND3, ATPase 8, and ATPase6, from NCBI (Table 1; details by individual in Table A1), increasing sampling density across Panama.

**Table 1:**
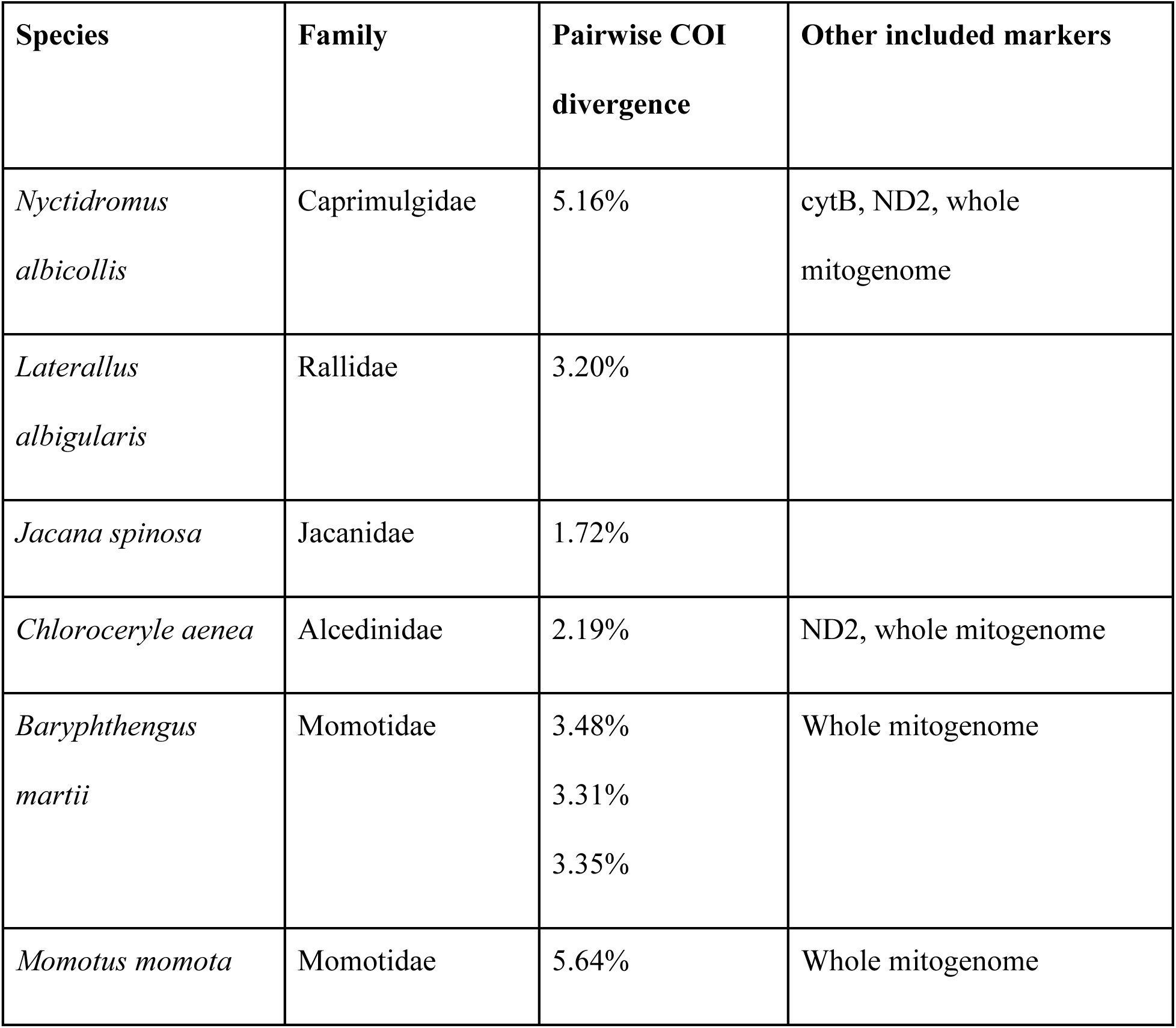
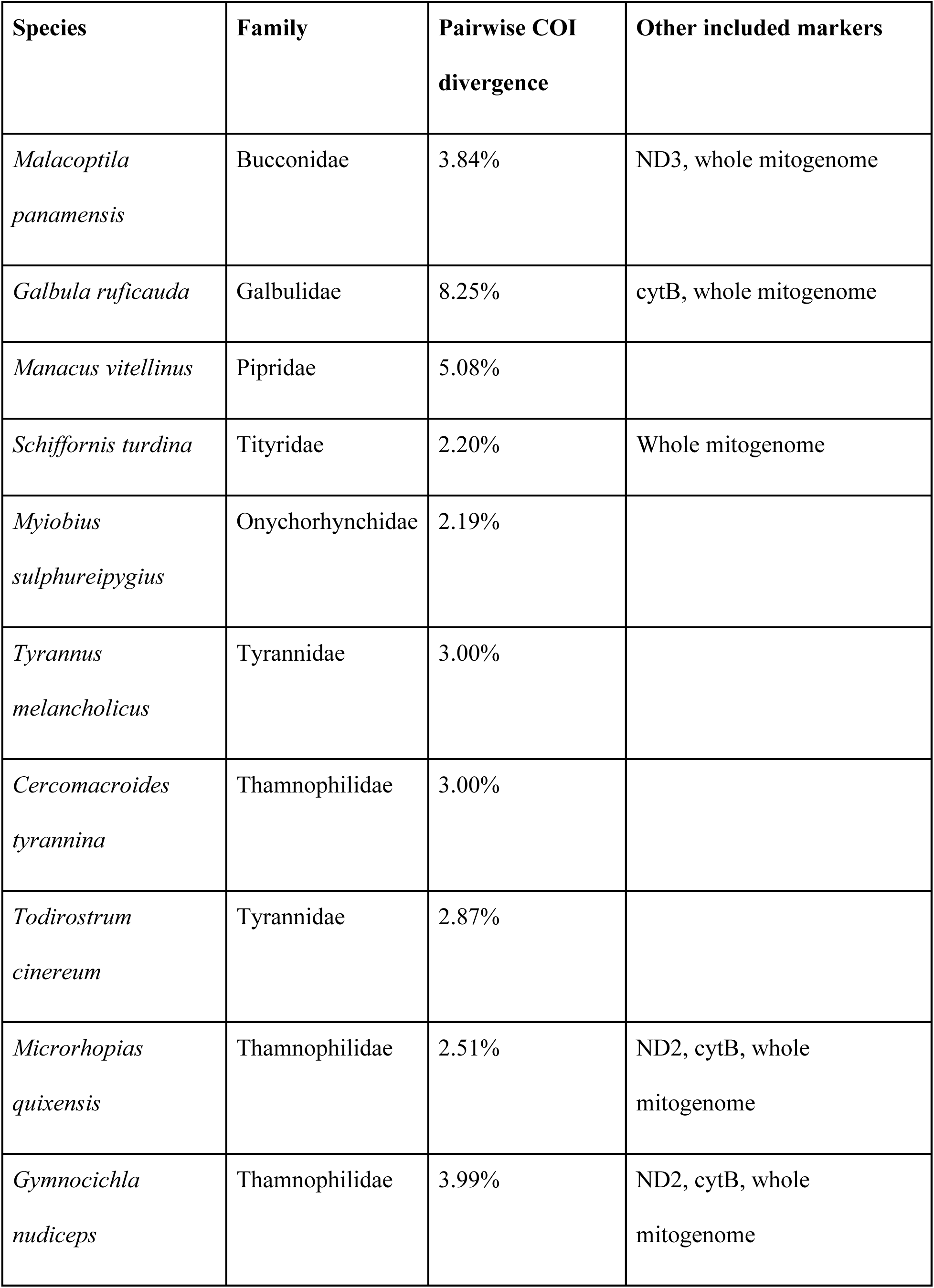
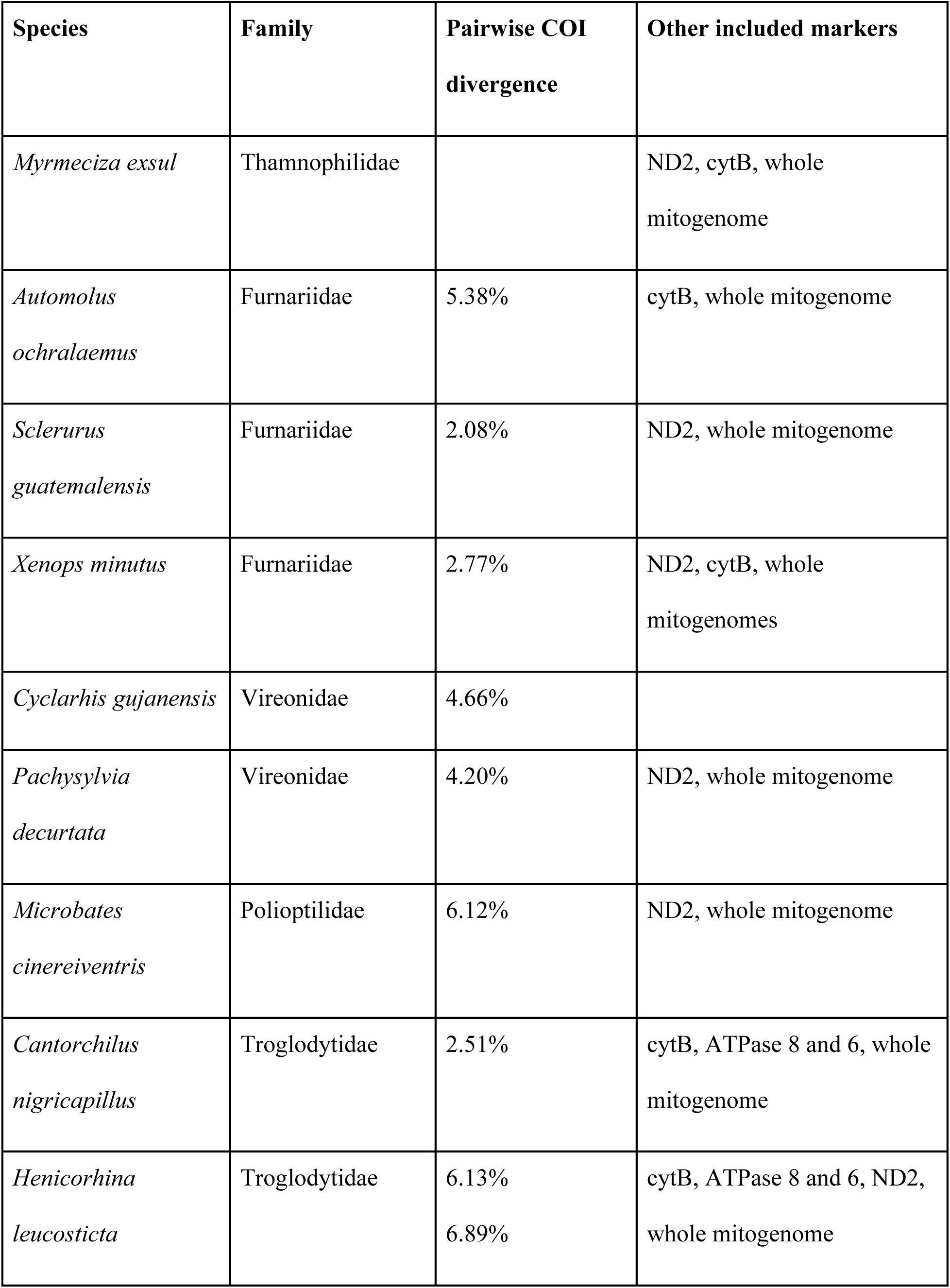
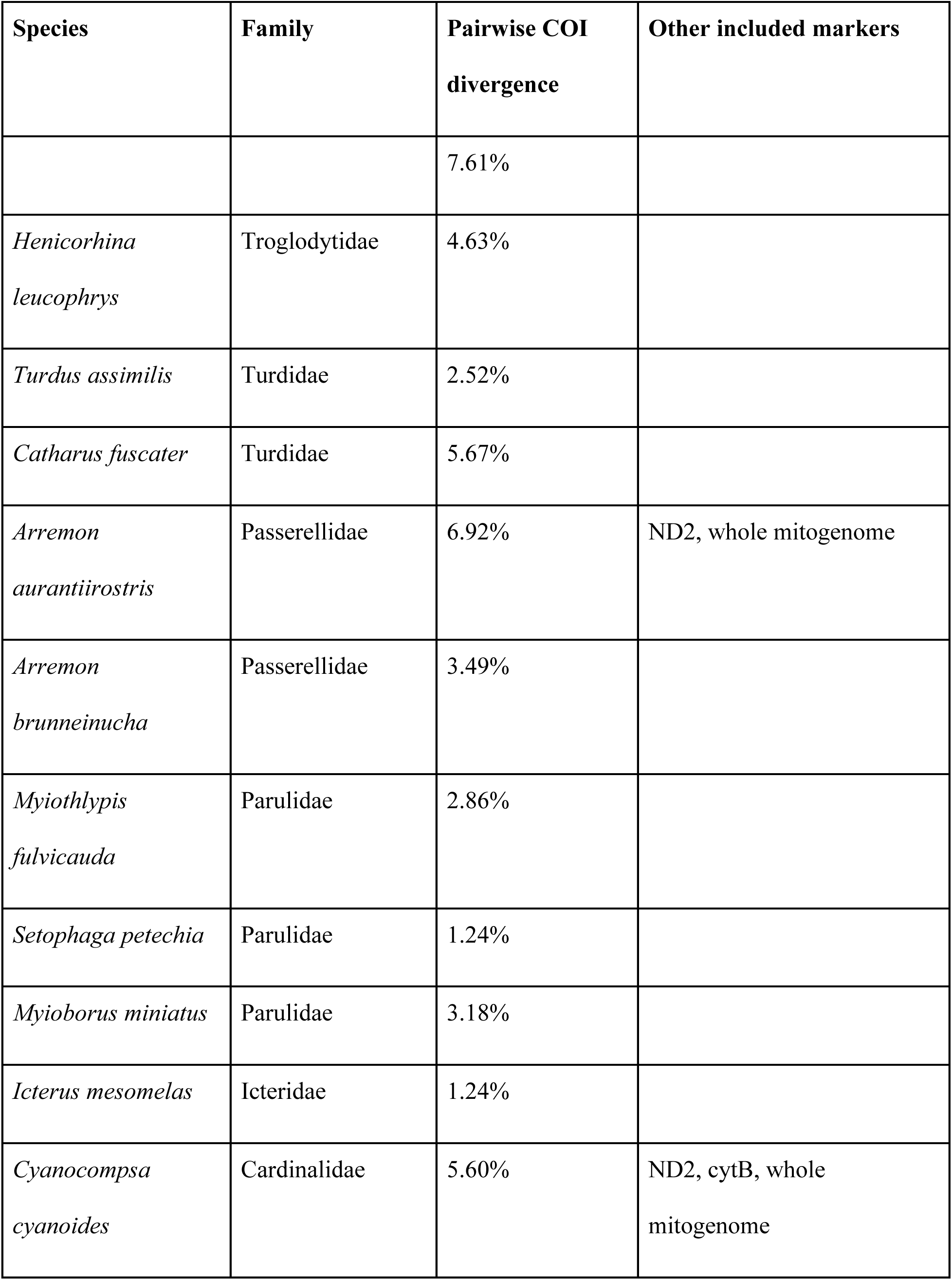
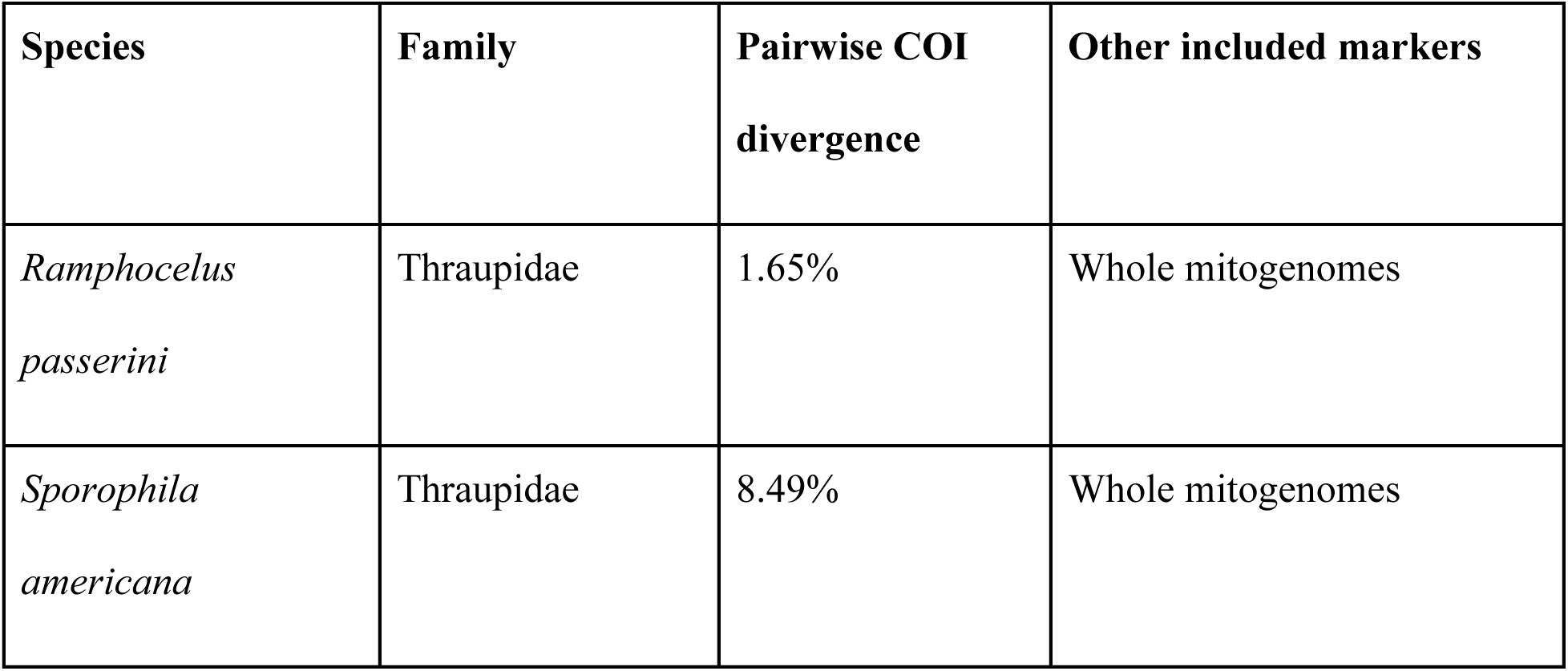
Evidence for cryptic divergence in Panamanian birds, organized by taxonomy. Results show species with COI splits defined as multiple BINs, with additional mitochondrial markers indicated. Distances calculated on BOLD-aligned sequences with K2P method. Species with more than two unique BIN assignments have pairwise COI divergence for all inter-BIN comparisons listed.

Assembly of one individual per population was performed using NovoPlasty v. 3.4 (Dierckxsens, Mardulyn, and Smits 2017), using either COI or ND2 as the seed, depending on availability. In some individuals with a high number of reads, we subsampled the initial reads with BBSplit, a part of the BBMap package (Bushnell 2014), to increase computational efficiency by only including putative mitochondrial reads in our assembly inputs. Mitogenomes were then aligned and annotated with MitoAnnotator (Iwasaki *et al*. 2013), from which we calculated the pairwise K2P genetic distance for each of the protein-coding genes between eastern and western populations.

### Quantifying Mitochondrial Splits

To estimate where mitochondrial splits (i.e., multiple BINs as defined by COI barcoding) occur, we generated locus-specific sequence alignments of available mitochondrial sequences for the 34 taxa above. Sequences were downloaded from BOLD and NCBI’s Genbank (Table A1). We then generated MUSCLE alignments (Edgar 2004) in MEGA7 (Kumar, Stecher, and Tamura 2016), using these to build neighbor-joining trees in PAUP* (Swofford 2001) or MEGA (Kumar, Stecher, and Tamura 2016) and ML trees in RaxML (Stamatakis 2014) using the GTR substitution model with Lewis ascertainment bias correction for 100 bootstrap replicates. For the 21 taxa with multiple mitochondrial loci available (Table 1), the BOLD barcodes were used to define groups, and then additional individual sequences were aligned to the whole mitochondrial genomes for that taxon, which had already been assigned to BINs, and haplotyped accordingly.

### Testing Predictors of Mitochondrial Divergence

We examined potential drivers of mitochondrial divergence using geographic, morphological, and ecological data. To test how ecological factors such as habitat openness, forest stratum, diet, and elevational distribution influenced the distribution of mitochondrial splits, we compiled and scored these data for all 659 resident, breeding landbirds of Panama, using species accounts from The Handbook of the Birds of the World Online (Billerman *et al*. 2020), supplementing individual species accounts as needed from Angehr and Dean (2010), Ridgely and Gwynne (1992), Stotz *et al*. (1996), and Wetmore (1965, 1968, 1972; 1984). Habitat was scored as forest, edge, or open (Table A2). Stratum was scored as ground (primarily terrestrial foraging, and/or prefers walking to flying), understory (forages primarily in undergrowth or directly above ground), midstory (primarily found in middle strata of forest, up to subcanopy), canopy (primarily found in the subcanopy and above), and aerial (forages above the forest canopy; almost exclusively swifts and swallows). We also included data on territoriality, hand-wing index (HWI), body size, and annual precipitation in range sourced from Sheard *et al* (2020).

Finally, we identified whether each species had been sampled across multiple geographic regions of Panama (Figure 1A), to identify the presence of geographic-based variation across the region. We tested for sampling biases in these three categories by comparing the total list and sampled subset by chi-squared tests to check if our 429 sampled taxa reflected the distribution of the above traits within the total Panamanian avifauna. Then, we tested whether those taxa which had been sampled across multiple geographic regions of Panama (Figure 1A) were representative of the total Panamanian avifauna. Taxa were considered widespread enough for inclusion in these tests if they occurred in two or more of the defined geographic regions of Panama (Figure 1A), which yielded a total of 181 species.

Of the 181 species in our study which had been widely sampled across Panama based on the above criteria, we then tested whether certain geographic, ecological, and morphological traits were disproportionately observed in species with two or more mitochondrial BINs. For each of the above ecological traits (stratum, territoriality, diet, habitat, HWI, body size, and mean annual precipitation), we tested using either a chi-squared test or student’s *t*-test whether there were significant differences in the representation of traits between split and non-split taxa.

### Testing for Coincident Timing of Splits

To assess if splits were coincident in time, and thus likely to have been driven by the same biogeographic events, we estimated divergence times in BEAST v. 2.6.2 (Bouckaert *et al*. 2014) as implemented on CIPRES (Miller, Pfeiffer, and Schwartz 2011). We generated an alignment of COI barcodes in MUSCLE, and trimmed ends to have no missing data. We used a strict clock rate of 1.8 % divergence per million years (Lavinia *et al*. 2016), gamma site model with JC69 substitution model (Naka and Brumfield 2018), and five fossil calibration points (Table A3). Fossil calibrations and the tree were given a log normal prior distribution. We ran this model for 1 billion MCMC generations, sampling every 10,000 generations, and assessed convergence (all ESS values > 200) in Tracer v. 1.7.1 (Rambaut *et al*. 2018). To generate a maximum clade credibility tree, we generated the MCC tree in LogCombiner with the first 20% as burn-in using median heights. We calculated the mean divergence time for each taxon using the rate of 1.8% divergence per million years previously found for avian COI (Lavinia *et al*. 2016), and compared this with the BEAST estimated means. With the 95% confidence intervals constructed in the latter, we then used these estimates to establish whether divergence times were broadly coincident.

## RESULTS

### Barcoding and Data Collection

We successfully barcoded 2,333 individuals from 429 species across Panama, 391 of which were resident landbird species (Table A1). This sampling was representative of Panamanian avifauna, as similar proportions of highland and lowland birds were present in the whole population of landbirds in Panama as in our sampled subset of species (!^2^ = 0, df= 1, *p*= 1.0) as well as similar proportions by diet (!^2^=13.34, df*=* 9, *p*=0.14) and habitat (!^2^=6.18, df*=* 3, *p*=0.10; Table 2). Of these 429 species, 181 (42.2%) were sampled across two or more geographic regions (Figure 1A), and were likewise representative of the whole population of resident landbirds (Table 2).

**Table 2:**
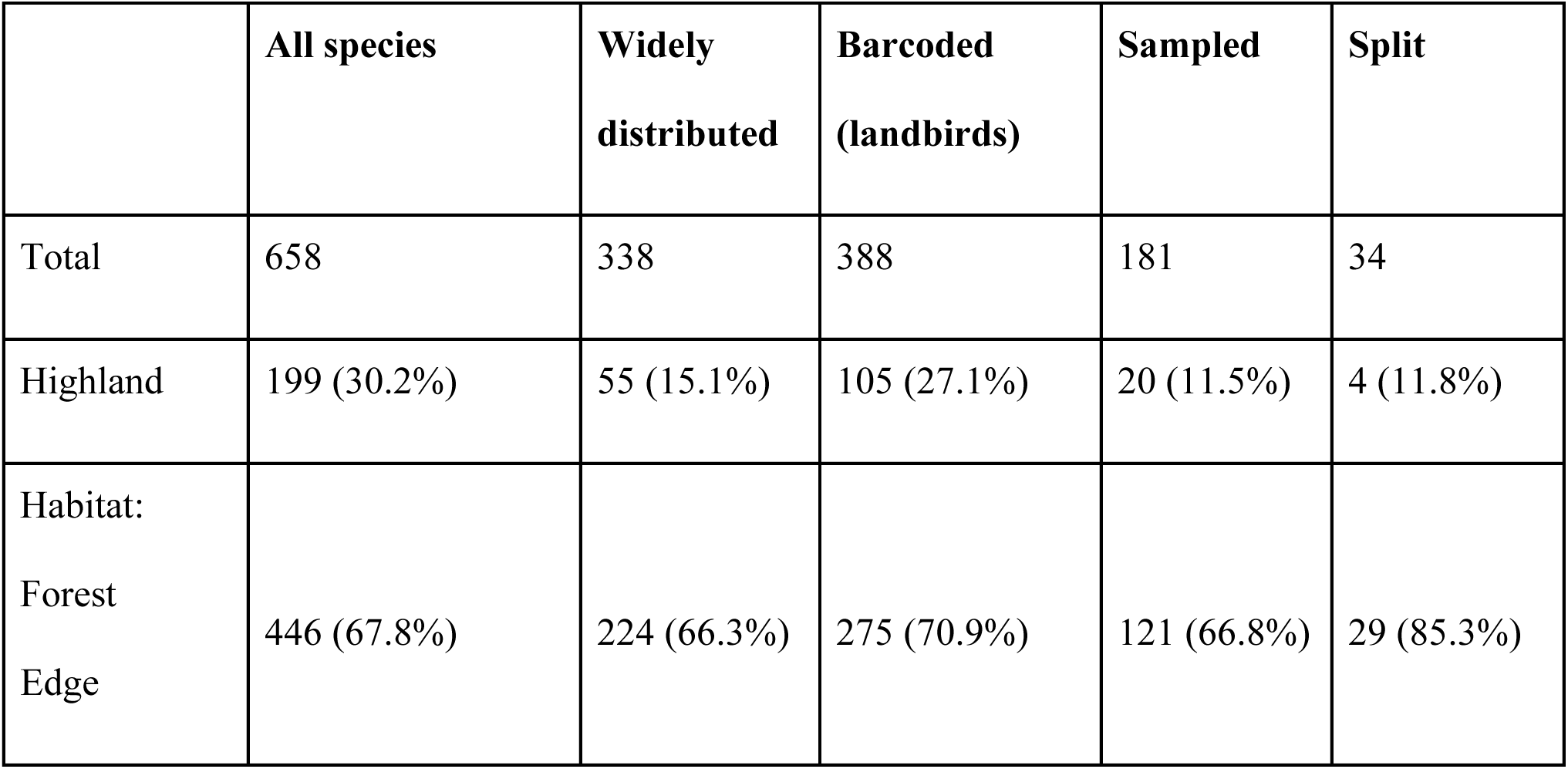
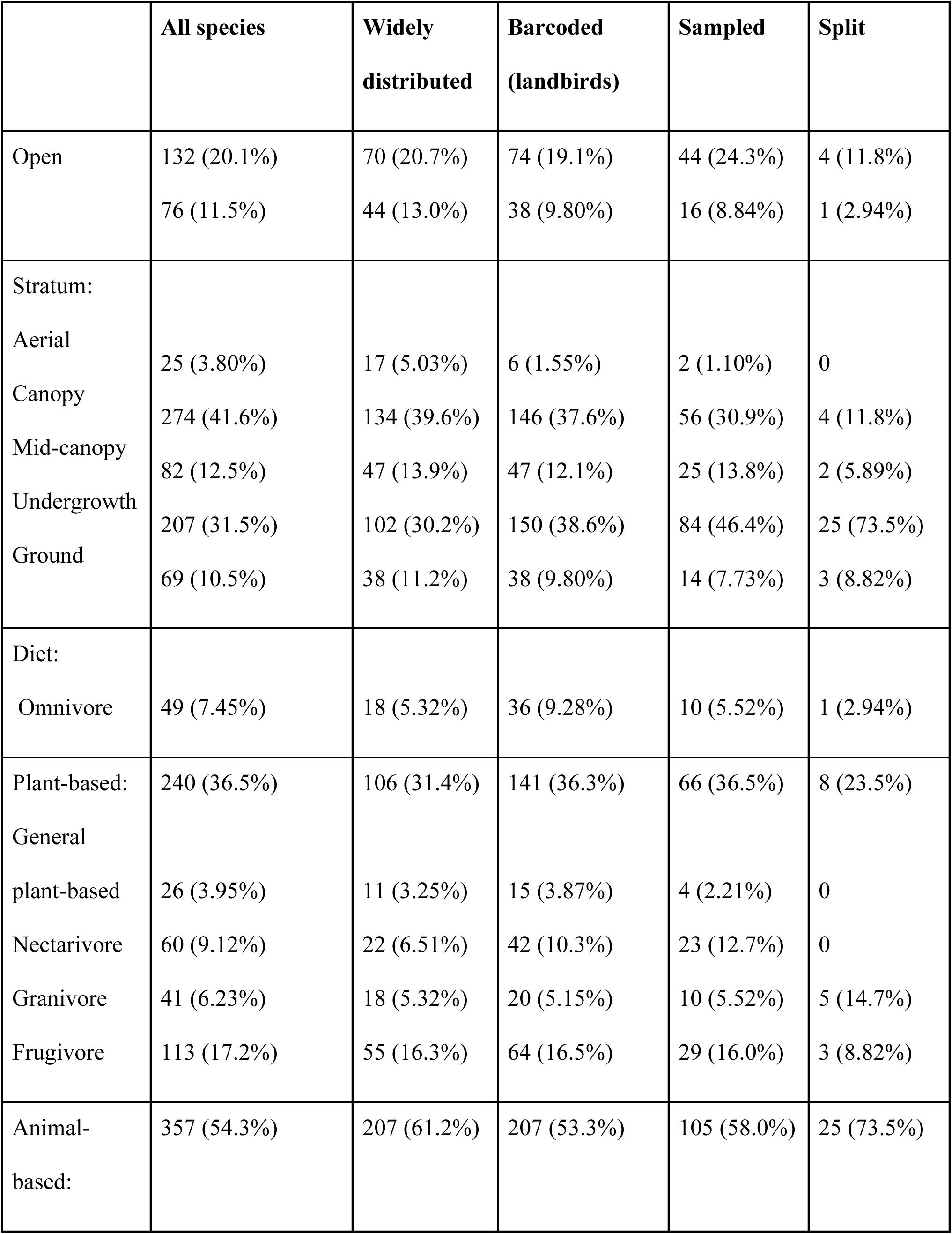
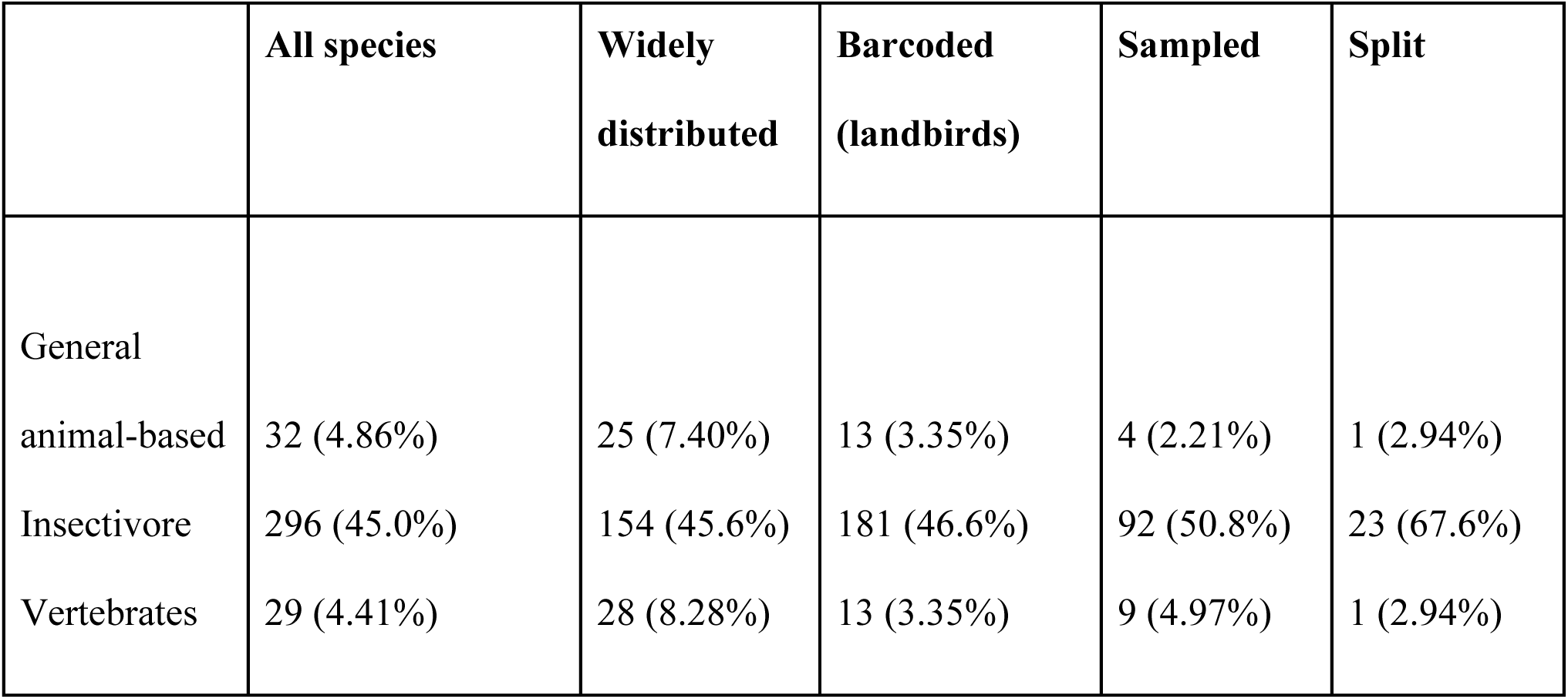
Correlates of cryptic divergence in Panamanian birds. Distribution of traits across species, showing across all 658 resident landbirds, those distributed across multiple regions, those barcoded (including both widespread and regional species), those resident landbirds with widespread distributions that were barcoded, and those within that last group found to have more than one BIN.

Thirty-four of these 181 taxa—18.8%— had more than 1 mitochondrial BIN, represented by a total of 419 individuals barcoded in BOLD. We increased this to a total of 634 individuals by adding 215 additional sequences from NCBI (Table A1). Twenty-one species were supplemented by this method, but the remainder did not have the required samples of each BIN’s whole mitochondrial genomes to allow the building of a multi-locus dataset.

### Mitochondrial Splits

Among landbirds, splits were observed in 20 of the 37 widely sampled families (Figure 2), accounting for 33% of the 61 resident landbird families documented in Panama. We characterized the geography of the 34 taxa with multiple BINs, plus two waterbirds (*Laterallus albigularis* and *Jacana spinosa*) not included in the prediction testing due to overall low sampling of waterbirds. Seven taxa, all lowland, had three BINs in Panama (Figure 3). Overall, we found 41 splits across 34 landbird species, out of the 181 taxa with sufficient sampling across multiple geographic regions.

**Figure 3:**
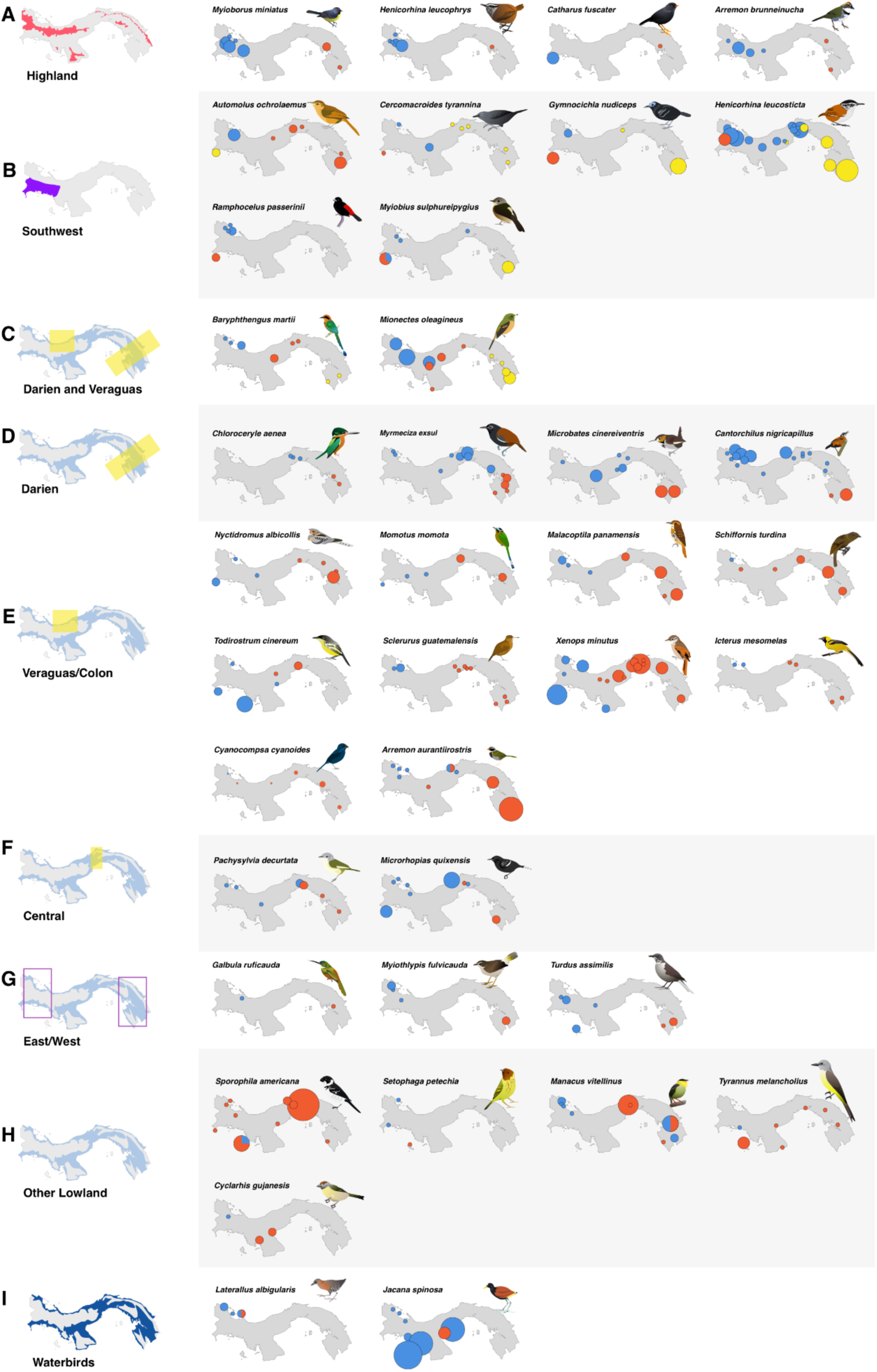
Geographic distribution of haplotypes in Panamanian birds. Haplotypes were defined initially by BIN for COI data, and then by sequence for additional markers for all taxa with observed mitochondrial breaks, grouped by geographic region of splits. Highland species (A) are separated from lowland birds, and lowland species are displayed by (B) southwest vs rest of Panama, with or without additional splits; (C) splits in both the Veraguas and Darién suture zones; (D) Darién suture zone splits; (E) Veraguas/Colón splits; (F) splits in central Panama, typically around Cerro Azul; (G) lowland taxa which have distinctive haplotypes in east and west, but lack sufficient sampling across the transect to determine the precise locality of the turnover; (H) taxa with idiosyncratic patterns that fit none of the above; and (I) waterbirds, which were generally excluded from analyses due to less consistent sampling. Dot colors indicate distinct BINs, size scaled by the number of samples from a given locality.

Geographically, we observed two primary patterns in the distribution of splits (Figure 3). The first, observed in six species, was a break between southwest Panama, in particular the Burica peninsula (Chiriquí province; Figure 3B), and the rest of Panama. This pattern was restricted to lowland taxa. The second pattern was one split between eastern and western Panama, observed in 35 splits (Figure 3D-G). This included both highland taxa (4 splits; Figure 3A) and lowland (31 splits; Figure 3D-F). In highland taxa, these splits were between the Cordillera Central and the highland areas of the east. However, in lowland taxa, there were two general clusters of regions of rapid geographic replacement of BINs across multiple taxa. The first, involving 15 splits (48% of lowland taxa with east-west splits), was along the Caribbean versant in Veraguas province, extending into Colón province in some cases (Figure 1B, 3E). The second, involving seven splits, was roughly located along the border of Darién and Panama provinces (Figure 1B, 3D). Three lowland splits were in central Panama (Figure 3F), and an additional three, representing distinct BINs in extreme eastern and western Panama, lacked samples which prevented us from locating the precise area of turnover (Figure 3G). One predicted geographic pattern that was not observed was differentiation of island and mainland taxa, despite island samples being included in most species and preferentially including taxa thought to represent distinct island groups (Table A1).

### Prediction Testing

While our overall dataset of 181 species may have been ecologically and geographically representative of resident landbird taxa (Table 2), the observed mitochondrial splits were associated with different factors. While insectivores made up 47% of non-split species, they accounted for significantly more (68%) of species with splits (!^2^=17.27, *df=* 7, *p*=0.02; Figure 4B). Forest birds, while comprising the majority of non-split species, at 62%, had even greater representation among the split species, at 85% (!^2^=6.49, *df=* 2, *p*=0.04; Figure 4C). When considering habitat stratum, we found that while non-split species were evenly distributed throughout strata, with only 40% being classed as understory residents, in split taxa understory birds were the overwhelming majority, at 74% of species (!^2^=14.04, *df=* 4, *p*=0.0072; Figure 4C). Hand-wing index (HWI) was significantly (*t=* -5.52, *df*= 154.29, *p*=<0.001) lower in split species (Figure 4A). Finally, split taxa were far more likely to be strongly territorial, with 62% of split taxa versus 35% of non-split (!^2^=12.04, *df=* 2, *p*=0.0024; Figure 4D).

**Figure 4:**
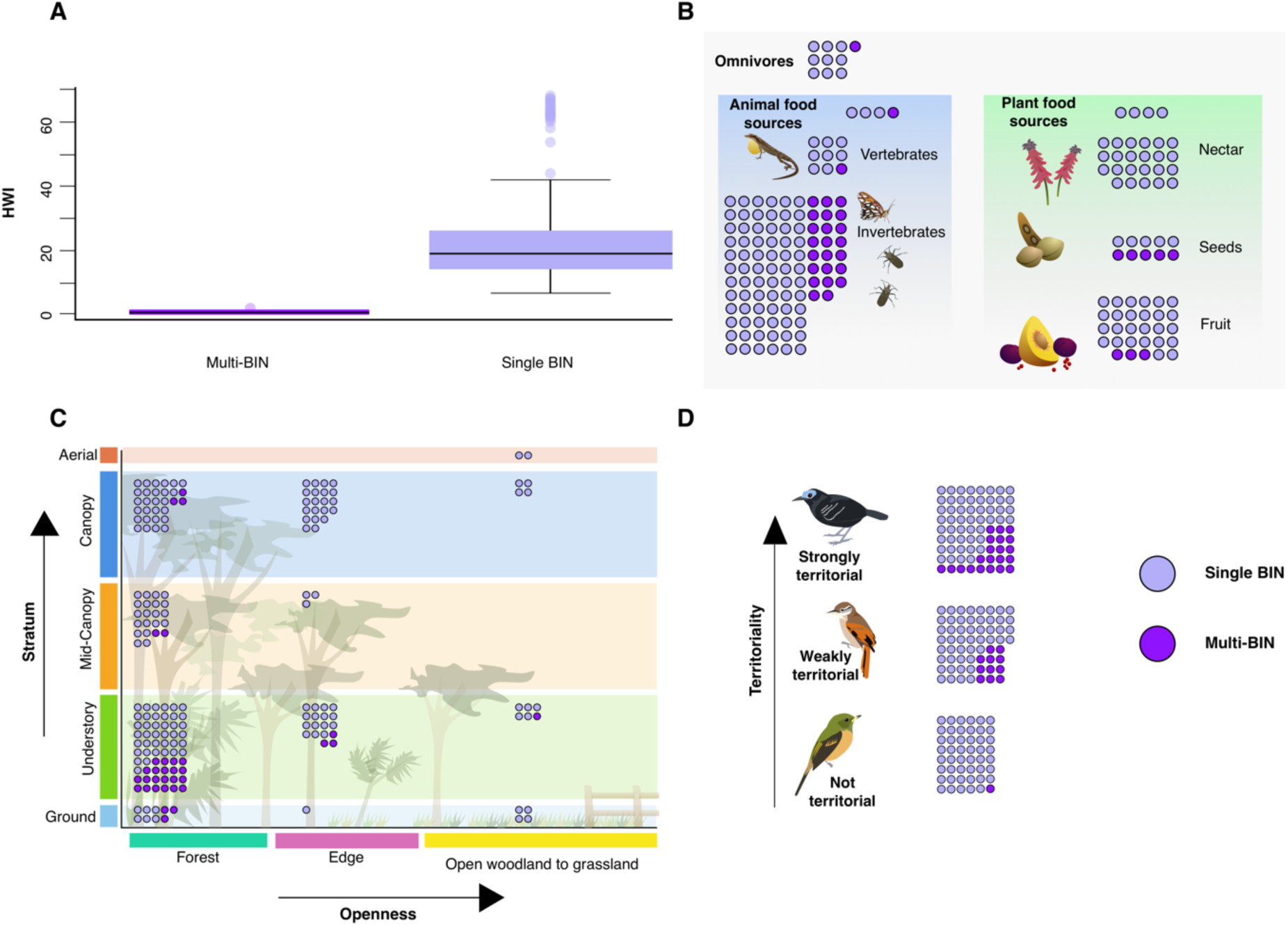
Associations of ecological traits with mitochondrial splits. Throughout the figure, dark purple circles represent taxa with two or more BINs, while light purple indicate those with only one. A) Hand wing index (HWI) is significantly lower in taxa with splits (*t=* -5.52, df*=* 154.29, *p*=<0.001), indicating lower physical dispersal capability is associated with mitochondrial turnover. B) Primary diet, showing that insectivores are represented significantly more than other dietary items (!^2^=17.27, df*=* 7, *p*=0.02) in taxa with splits. C) A visual representation of habitat use, showing that habitat type as measured by openness (!^2^=6.488, df*=* 2, *p*=0.039) and stratum (!^2^=14.04, df*=* 4, *p*=0.007) are associated with mitochondrial turnover, with it becoming increasingly likely in the closed forest understory. D) Despite the relatively even distribution of territoriality across our sample, strongly territorial taxa were overrepresented among those with mitochondrial turnover (!^2^=12.04, df*=* 2, *p*=0.002).

Some traits, however, were represented at similar proportions in both split and non-split species (Table A2). Highland taxa were a minority of species in both cases, at 11% of non-split and 12% of split species (!^2^=4.10 × 10^-29^, *df=* 1, *p*=1). Habitat mean annual precipitation was similar for both, at 2209 mm/yr in split and 2179 mm/yr in non-split (*t=* 0.28, *df*= 48.22, *p*=0.78). Finally, scaled body size was largely similar, with non-split species being very slightly larger, but not significantly so (*t=* 0, *df*= 292, *p*=1).

### Timing of Splits

Depths of splits varied considerably. Pairwise differences in COI ranged from 1.24% to 8.49% for those taxa defined as having multiple BINs by BOLD. Median pairwise divergence was 3.31%. These are equivalent to between 689 kya and 4.71 mya (Figure 5), with a median of 1.84 mya. However, estimates from BEAST were substantially older than from pairwise estimates, as expected with dates from coalescent methods, and our confidence intervals were frequently very wide (Figure 5) and do not coincide temporally.

**Figure 5:**
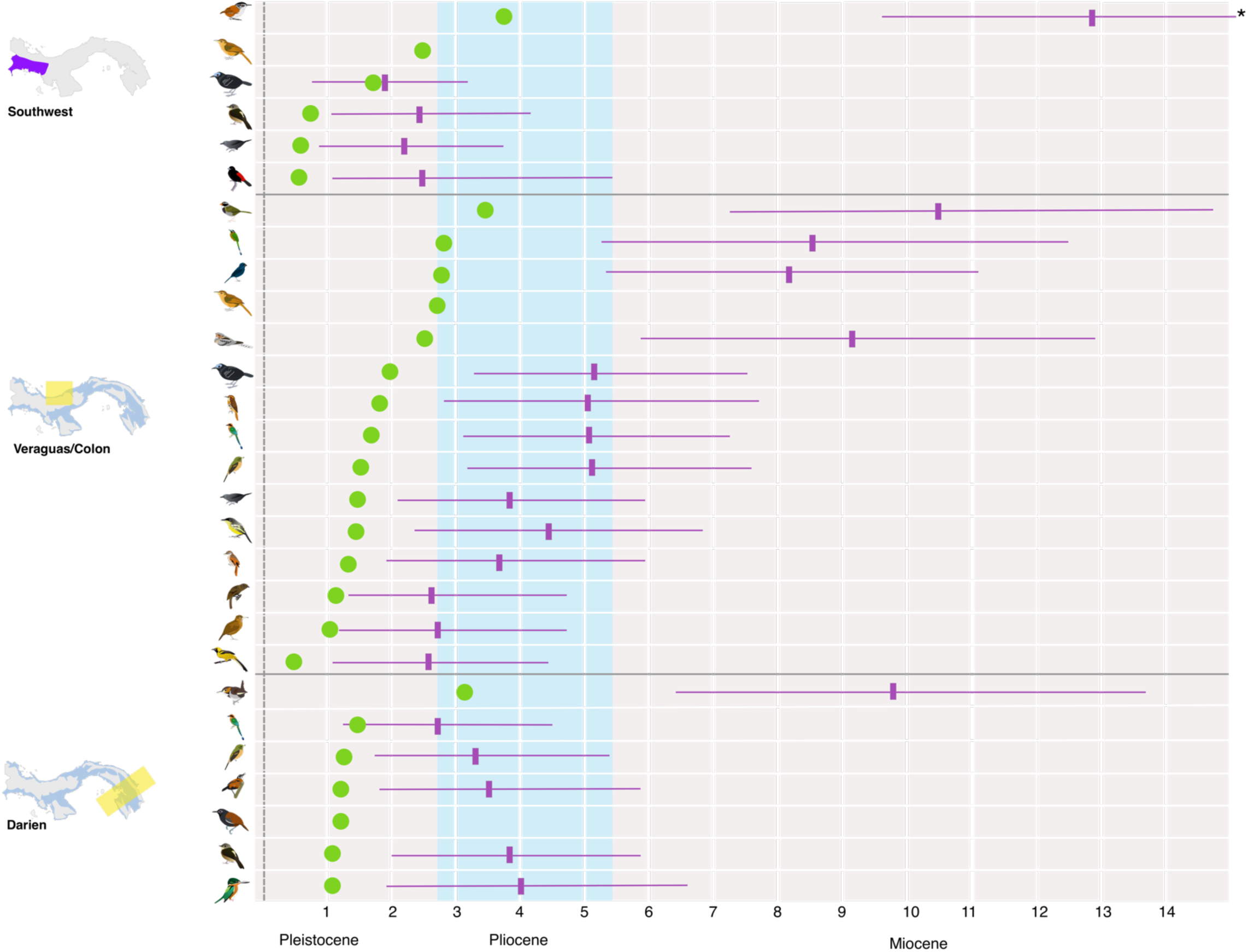
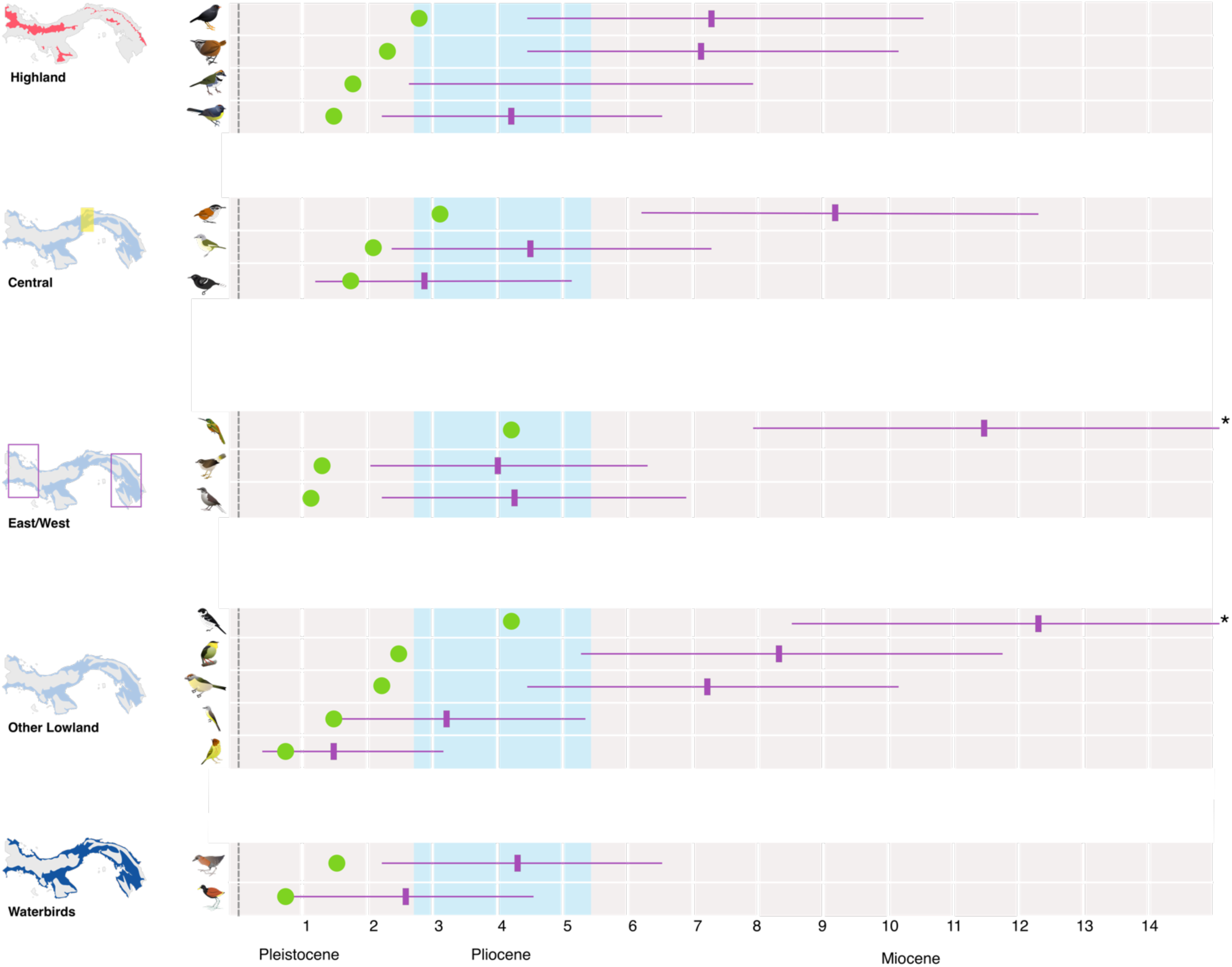
Timing of mitochondrial splits for Panamanian birds in millions of years. Time was calculated from pairwise COI divergence (green) and in BEAST2 (purple, shown with median and 95% confidence intervals [CIs]). Splits are grouped by region as in Figure 3, with those taxa with multiple splits being shown for each. Some CIs are truncated due to space (indicated with asterisk); see Table 3 for full details.

**Table 3:**
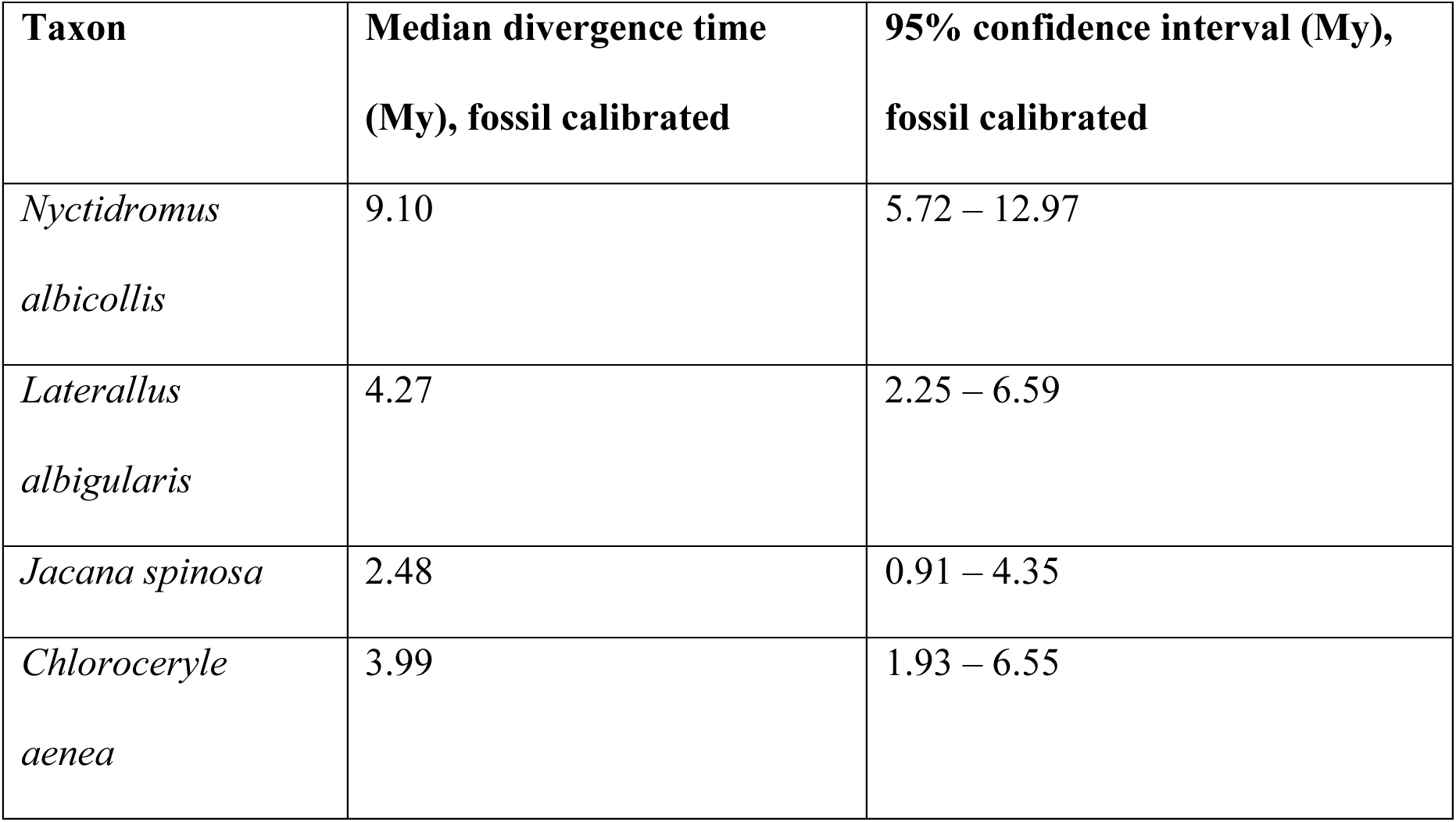
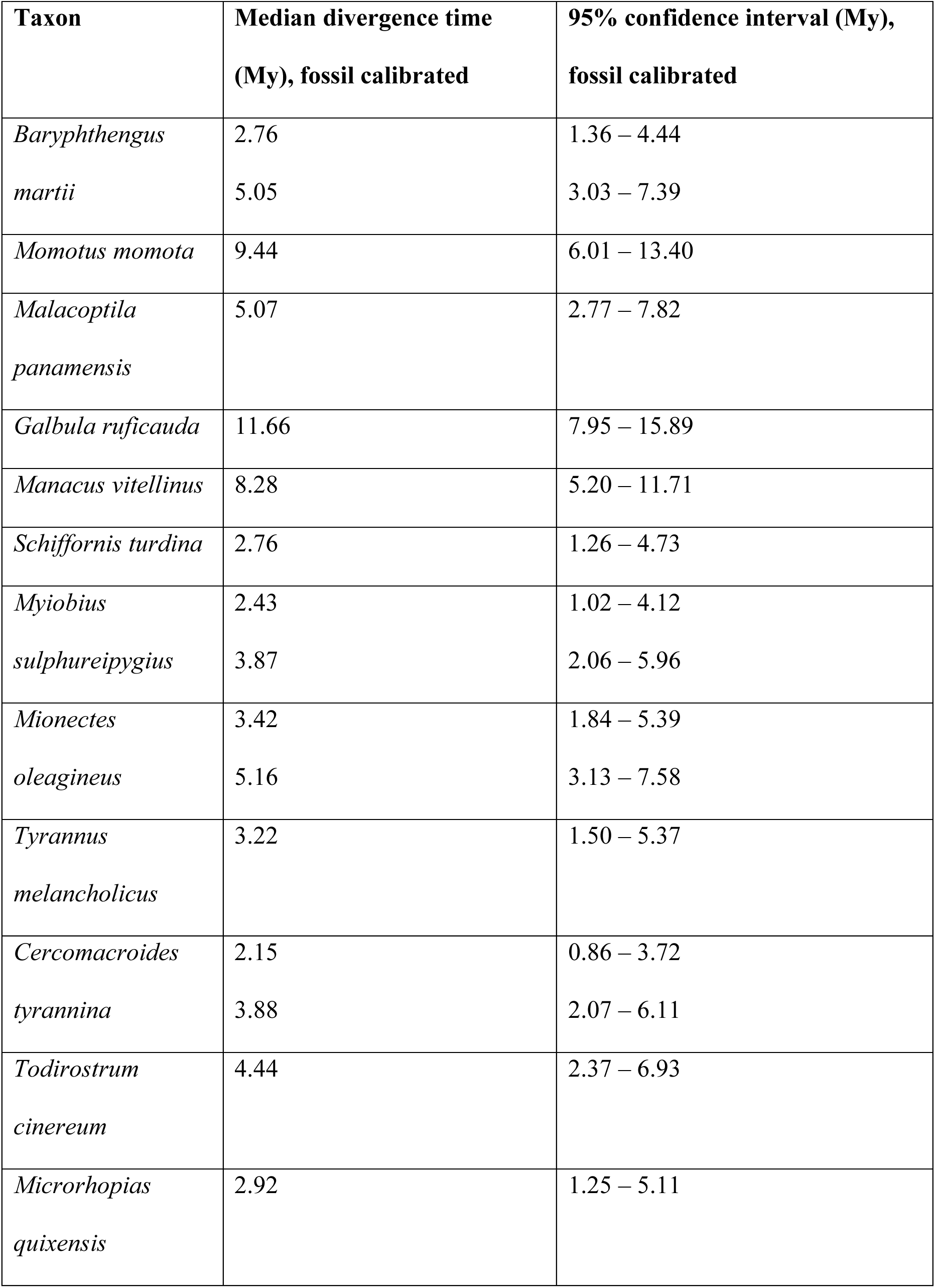
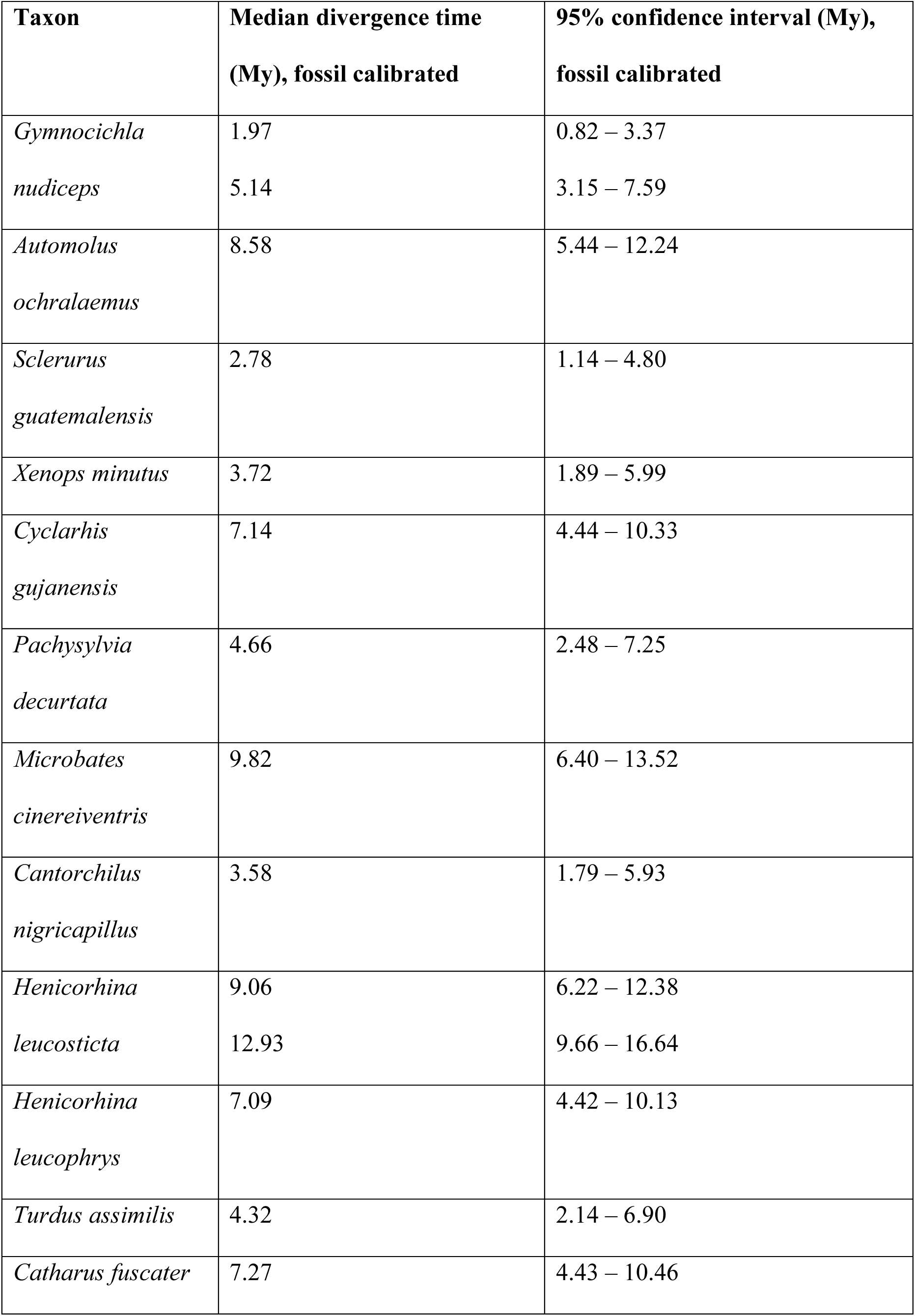
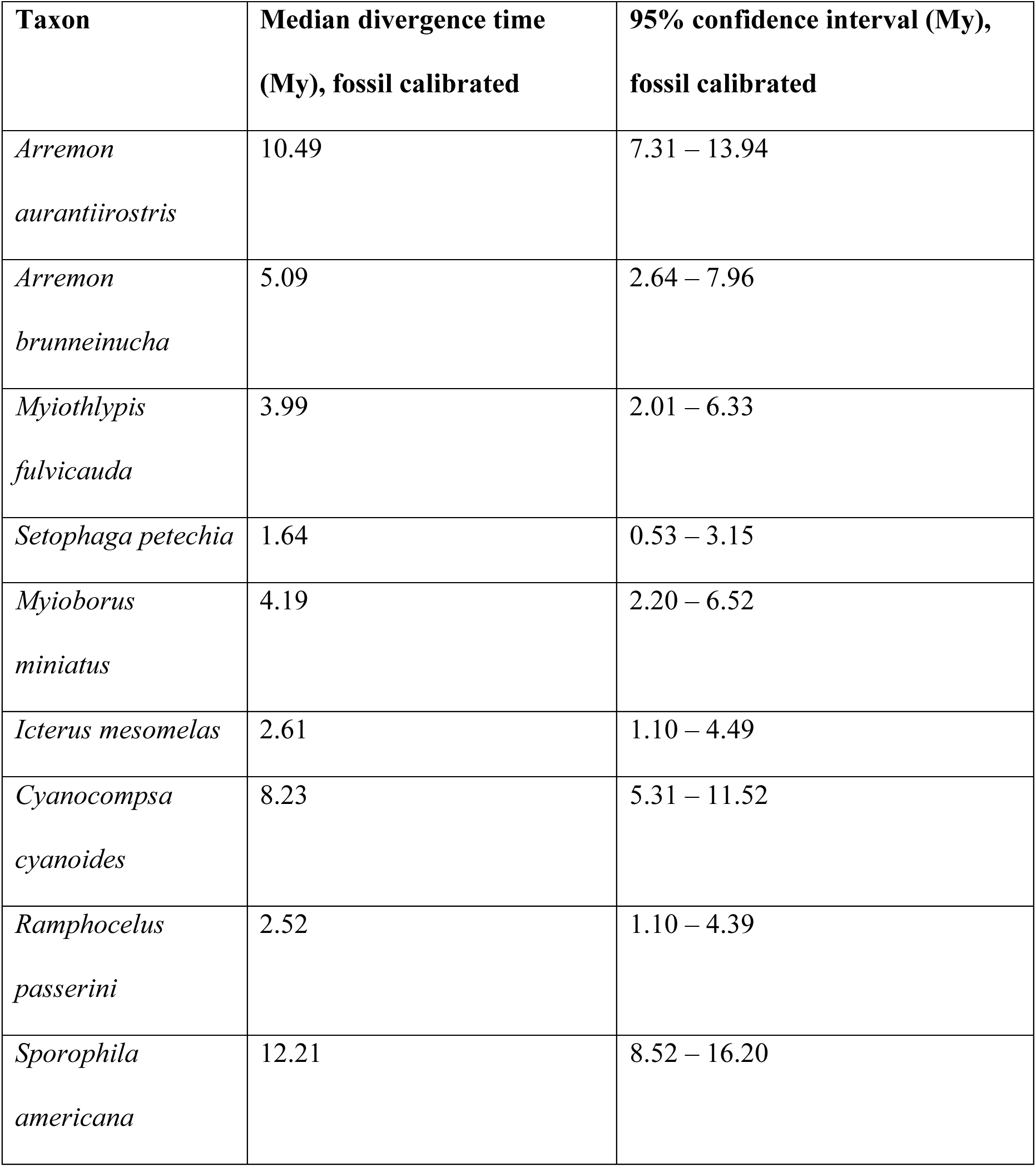
Estimated time to the most recent common ancestor in Panamanian birds. Estimates were calculated in BEAST2 for the 34 taxa with multiple BINs. For taxa with three BINs, estimates for both split events are provided.

## DISCUSSION

Using mitochondrial barcoding, we both identified substantial cryptic diversity in Panamanian birds and found support for ecological drivers of diversification in the region. We found that the frequency of cryptic diversity in Panamanian resident landbirds is 18.8% (Table 2). This is higher than estimates from similar barcoding efforts in other geographic regions, including the 2.7% of North American birds (Kerr *et al*. 2007), 11% of Korean birds (Yoo *et al*. 2006), 7.5% of Palearctic birds (Kerr *et al*. 2009), 3.3% of Argentinian birds (Kerr *et al*. 2009), and 3.6% of South American birds more generally (Tavares *et al*. 2011). Due to variations in study designs, however, some caution is needed in directly comparing these without controlling for differences in sample size, taxonomic biases, and other potential sources of variation. While other studies have found higher incidence of potential cryptic diversity, these tended to focus on narrow subsets of birds that may be particularly predisposed towards splits. For example, Milá *et al*. (2012) found evidence of interspecific-level variation within 33 of 40 forest understory birds in the Amazon, a habitat profile that we find to be significantly overrepresented in lineages with potential cryptic species in our study. In general, many barcoding studies have tended to focus on large-scale questions of overall divergence, rather than explicitly examining whether specific ecological traits were over-or under-represented in taxa with potential splits (Yoo *et al*. 2006; Kerr *et al*. 2007, 2009; Kerr *et al*. 2009; Tavares *et al*. 2011). Even in light of these differences in study design and underlying questions, we find a notably high rate of cryptic diversity in Panamanian birds compared with other studies.

### Geographical Patterns of Cryptic Diversity

We sampled widely across Panama and evaluated several previously proposed biogeographic patterns in cryptic diversity (Wetmore 1959; Summers *et al*. 1997; Anderson and Handley 2001; Miller *et al*. 2011; Kaviar, Shockey, and Sundberg 2012; Miller *et al*. 2015). The most common geographic pattern that we observed was differentiation between eastern and western Panama. These splits could be further subdivided into three general patterns: disjunct highland populations from the Cordillera Central and the highlands of eastern Panama, a suture zone in the Caribbean versant of Veraguas and Colón, and a second suture zone in Darién (Figure 1B).

Many of the taxa included samples from the various islands of Panama, including Isla Coiba in the Pacific and the Caribbean islands of San Cristobal, Bastimentos, Cayo Agua, and Escudo de Veraguas. Islands are a natural focus of investigation for undescribed biodiversity, and previous studies have found evidence for island endemics in both birds and other taxa in Panama (Summers *et al*. 1997; Anderson and Handley 2001; Kaviar, Shockey, and Sundberg 2012). Escudo, for example, has had four of its eight to ten resident breeding birds described as endemic subspecies (Wetmore 1959). The Escudo hummingbird (*Amazilia (tzacatl) handleyi*) is both phenotypically and genetically distinct from mainland populations (Miller *et al*. 2011) and is sometimes treated as a separate species (Wetmore 1968; Angehr and Dean 2010). However, overall, we found low mitochondrial divergence between mainland and island taxa. This is likely due to their close proximity to the mainland, such as in the case of the islands of Bocas del Toro (approximately 15 km for the furthest islands of Bocas del Toro, and 20 km to Isla Escudo) and Coiba (24 km to mainland) which makes it likely that they were intermittently connected to the mainland during Pleistocene sea-level fluctuations (Miller *et al*. 2011). Escudo, the furthest of the Caribbean islands, is estimated to have become isolated only around 9000 years ago (Summers *et al*. 1997; Miller *et al*. 2011). This lowers the probability of the development of distinctive island endemics (MacArthur and Wilson 1963; Mayr 1965). Furthermore, the majority of island avifauna are also found on the mainland (Wetmore 1965, 1968, 1972; Wetmore, Pasquier, and Olson 1984; Ridgely and Gwynne 1989; Angher and Dean 2010), suggesting that there has been a high degree of connectivity over time. Indeed, *Setophaga petechia* was the only taxon with a distinctive BIN on Isla Coiba, but we suspect this was likely an individual of the migratory subspecies (included unintentionally, as we overall excluded migratory taxa) rather than the resident, as both occur in Panama. While island populations may be a source of haplotype diversity in some taxa (González *et al*. 2003), they may not be sufficiently diverged to be split under the BIN system.

An area where we expected to find unrecognized diversity is the Pacific coast of Chiriquí, particularly the Burica Peninsula (Figure 1A). This region, separated from most of the rest of Panama by the Cordillera Central, receives markedly less precipitation than the rest of the country (Wetmore 1965), and is thus home to much more xeric ecosystems than on the rainy Caribbean side (Blanco *et al*. 2013). We found support for this region being a hotspot of unrecognized avian diversity, with six of the 41 splits located in this relatively small area (Figure 3).

The four highland splits are all between the two main highland regions of Panama (Figure 4). The predominant east-west split is consistent with the numerous species-pairs previously observed to follow this pattern (Wetmore 1965, 1968, 1972; Wetmore, Pasquier, and Olson 1984; Ridgely and Gwynne 1992; Angehr and Dean 2010). Likewise, the Azuero Peninsula has been previously noted for several potential endemic species (Miller *et al*. 2015). However, across our overall dataset, highland splits are very much in the minority of those observed, making up only 14.7% of splits.

### The Lowlands are a Hotspot for Cryptic Diversity

We found a disproportionate number of lowland taxa with cryptic species-level variation. This is likely due to the fact that previous efforts have focused on identifying potential cryptic species in taxa with disjunct ranges such as highland species. Indeed, highland species showed splits of equivalent depth, but have likely already been defined as separate species (e.g., *Basileuturus ignotus,* Todd 1929; *Scytalopus,* Krabbe and Schulenberg 1997, Cadena *et al*. 2020; *Tangara dowii* and *T. fucosa,* Burns and Naoki 2004). Thus, few highland taxa met our geographic sampling criteria of being present in two or more regions (Figure 1A), and they represented only 15.1% of widely distributed species and 11.5% of species barcoded across multiple geographic regions, despite making up more than 30% of the total avifauna (Table 2). This discrepancy in the rates of splits in lowland and highland taxa is potentially due to bias from prior taxonomic description of variation. While this bias may partially explain the discrepancy, it does not explain what may generate the lowland variation we observed.

While many of Panama’s lowland bird species are continuously distributed (Wetmore 1965, 1968, 1972; Wetmore, Pasquier, and Olson 1984; Ridgely and Gwynne 1992; Angehr and Dean 2010), this may not have historically been the case, as Pleistocene climate variation may have caused ranges to contract (Haffer 1969; Moritz *et al*. 2000; Weir, Bermingham, and Schluter 2009; Smith, Amei, and Klicka 2012; Lavinia *et al*. 2015). Most of the split bird species (85%) are forest birds (Table 2), and the increased aridity during the Pleistocene is thought to have caused widespread advancement of savannah into formerly forested areas throughout the tropics (Haffer 1985; Webb 1991). Thus, currently continuously-distributed species might not have always been connected, and this climatic history may have driven the diversification of taxa across Panama. However, it is unclear the extent of forest contraction in Panama throughout the Pleistocene (Bush and Colinvaux 1990; Colinvaux, De Oliveira, and Bush 2000), and we find little support for the observed divergence dates coinciding to even roughly similar time periods (Table 1).

Clearly, while Panama’s diverse landscape has played a role in generating its rich avian diversity, this temporal incongruence of divergences (Figure 5) suggests that biogeographic models centering geological history are not the only drivers of cryptic diversification in the region. Previous work in Panama and lower Central America (LCA) more broadly has focused on the timing of the emergence of the isthmus (Smith and Klicka 2010), and in the resulting biotic interchange between North and South America (DaCosta and Klicka 2008; Weir, Bermingham, and Schluter 2009). However, it is unlikely that dispersal following the formation of the isthmus is responsible for the patterns of turnovers we see in our taxa. Our oldest splits (∼4 mya) predate the 3 mya date generally accepted for the formation of the isthmus (Leigh, O’Dea, and Vermeij 2014a), while the youngest (< 1 mya) are well after this time. However, the gradual emergence of the isthmus would possibly better explain some of the splits (O’Dea *et al*. 2016). In particular, we find a group of splits in the Veraguas/Colón suture zone around the Pleistocene-Pliocene boundary (Figure 5), which line up with a potential episode of seawater breaching the newly formed isthmus approximately 2.45 mya (Groeneveld *et al*. 2014; O’Dea *et al*. 2016). Though overall our dates do not support a scenario of dispersal prior to the formation of the isthmus, followed by secondary contact in most lowland cases, a more gradual process in which a changing landscape of newly emerged land acted as a tenuous bridge for dispersal over a longer period may be possible.

### Ecological predictors of mitochondrial turnover

If biogeographic factors alone cannot explain the generation of diversity in lowland Panamanian birds, what other factors may be important? Our results highlight how ecological traits alone have the potential to drive divergence, even in a landscape that has historically lacked obvious barriers to dispersal (Bush and Colinvaux 1990). Specifically, we found that traits such as habitat use, diet, and morphology can be effective in generating diversity, with the majority of taxa with mitochondrial splits sharing specific ecological traits. Insectivores (68% of sampled taxa with splits vs. 47% non-split taxa), forest birds (85% vs 62%), understory foragers (74% vs 40%), and strongly territorial species (62% vs. 35%) were all overrepresented in lineages with mitochondrial turnover (Figure 4).

A potential explanation for at least some of the role of these ecological factors is detectability. Detectability can have large ramifications for studying groups in difficult-to-reach areas, particularly in the context of conservation (Clark and May 2002; Ducatez and Lefebvre 2014; McKenzie and Robertson 2015; Smith *et al*. 2020). This can also extend to taxonomic considerations, as more easily observed and “charismatic” organisms are more likely to be oversplit (Pillon and Chase 2007). In particular, some habitats and strata are more likely to harbor cryptic species not because they are inherently more likely to create diversity, but simply because they are more difficult to observe. Therefore, the apparent overabundance of cryptic types in understory forest birds may be a relative scarcity of human observations. However, the overall well-described Panamanian avifauna make a lack of observation less likely than elsewhere in the Neotropics (Angehr and Dean 2010; Ridgely and Gwynne 1992; Wetmore 1965, 1968, 1972; Wetmore, Pasquier, and Olson 1984), and many splits within inconspicuous understory birds have been previously described (e.g. Saucier, Sánchez, and Carling 2015).

The traits observed to have a strong link to mitochondrial divergence are all associated with low dispersal capability. This is most obvious in the much lower HWI of birds with mitochondrial splits. HWI is a well-recognized proxy of dispersal ability (Claramunt *et al*. 2012; Claramunt and Wright 2017; Sheard *et al*. 2020), as it describes wing shape and thus the ability for sustained flight (Kipp 1959; Lockwood, Swaddle, and Rayner 1998; White 2016; Claramunt and Wright 2017; Sheard *et al*. 2020), and the lower HWI of split species shows that lower dispersal ability is significantly associated with mitochondrial turnover. The association of ecological factors linked to lower dispersal ability with mitochondrial turnover holds through other tested traits. Forest birds have much lower dispersal abilities than edge or open area species (Moore *et al*. 2008; Burney and Brumfield 2009; Weir, Bermingham, and Schluter 2009), especially those which primarily use the understory (Burney and Brumfield 2009; Woltmann and Sherry 2011). Likewise, strongly territorial species were overrepresented in the split taxa. As these species are less likely to disperse once they have established a territory (Greenwood 1980), this provides further weight to the role of dispersal.

Our finding of insectivores having greater frequency of splits may also be tied to differences in dispersal ability. Previous studies have found strong evidence that diet type, and especially the extent to which a given species relies on plant-based food sources, can shape dispersal and demography (Westcott and Graham 2000; Moore *et al*. 2008; Burney and Brumfield 2009; Miller *et al*. 2021). While both plant and animal food sources are typically available year-round in the tropics, seed, nectar, and fruit tend to be spatially and temporally clustered (Morton 1973; Levey and Stiles 1994). While insectivores may reliably find arthropods in a given home range (Levey and Stiles 1992; Burney and Brumfield 2009), birds feeding primarily on fruit, seeds, and nectar may need to travel more widely to seek out food sources throughout and between years (Westcott and Graham 2000). Furthermore, the relative availability of these resources varies between years to different extents. While arthropods are certainly subject to population cycles, they are usually less extreme (Jahn *et al*. 2010) than the fluctuations between mast years and lean years in fruit and seed-bearing species typically relied on for food by frugivorous and granivorous birds (Faaborg, Arendt, and Kaiser 1984; Wheelwright 1986; Levey, Moermond, and Denslow 1994; Brawn, Karr, and Nichols 1995; Ryder and Sillett 2016; Macario *et al*. 2017). As a result, birds which primarily feed on plants are more subject to boom-and-bust population dynamics (Faaborg, Arendt, and Kaiser 1984; Greenberg and Gradwohl 1986; Şekercioğlu *et al*. 2002; Woltmann and Sherry 2011; Sherry *et al*. 2020), and during boom years will experience increased dispersal, potentially connecting populations more regularly and slowing the accumulation of divergence between them.

The importance of traits directly or indirectly tied to dispersal ability may be to be driving a simple isolation-by-distance (IBD) effect. Poor dispersers will develop greater divergence across a given space than better dispersers, so that further populations will be increasingly genetically differentiated (Wright 1943, 1946; Slatkin 1993). However, while that may play a part for some of the taxa in our study, it is unlikely for all the observed splits. While some taxa, such as *Mionectes oleagineus* and *Baryphthengus martii*, have repeated mitochondrial breaks with increasing divergence across Panama (Figure 4), others have sharp turnovers within Panama with equal or greater divergence estimates, yet these haplotypes are still found hundreds of kilometers away in Nicaragua (*Arremon auraniirostris*, *Cyanocompsa cyanoides*), Honduras (*Arremon aurantiirostris*), Belize (*Cyanocompsa cyanoides*), and Ecuador (*Cantorchilus nigricapillus*). Thus, it is likely that dispersal is the driver of divergence in concert with other ecological factors in many cases.

Despite its small size, Panama is home to remarkable avian diversity, with 1000 currently recognized species occurring in the region (Wetmore 1965, 1968, 1972; Wetmore, Pasquier, and Olson 1984; Ridgely and Gwynne 1992; Angehr and Dean 2010). While some of this is likely due to the region’s position as a literal bridge between North and South America (DaCosta and Klicka 2008; Smith and Klicka 2010; Leigh, O’Dea, and Vermeij 2014b), the region itself plays an important role in generating biodiversity. The diverse elevational and climatic range of habitats across the isthmus provide opportunity for biogeographic scenarios leading to endemism (Stiles 1983; Barrantes 2009; Chavarría-Pizarro *et al*. 2010; Batista *et al*. 2020), but this alone may not be the only driving force behind avian diversity. Consistent with other recent work in the region (Miller *et al*. 2021), we find that ecological factors are strongly associated with cryptic diversity, particularly in lowland Panama, emphasizing the need to look beyond physical barriers as drivers of Neotropical biodiversity.

## CONCLUSIONS

Panama is widely recognized as an area of high biodiversity; however, we find that avian diversity is substantially underestimated. Potential cryptic species are in some cases associated with landscape and geography, such as highland taxa and those in southwestern Chiriquí, but the bulk of the observed splits are in lowland taxa in the absence of geographic barriers. The varying ages of the observed splits, from approximately 0.75–4.2 mya, makes it unlikely that all the observed variation is driven by a single historical factor. We find instead strong correlations between dispersal ability, both directly (HWI) and indirectly (through ecological traits such as habitat, diet, and territoriality), and the occurrence of mitochondrial turnover. This suggests that intrinsic ecological and life history traits can be a major factor in driving species turnover and the accumulation of biodiversity in the tropics. This also illustrates how examining cryptic species can provide insights into the ecological and evolutionary processes that shape this diversity. The potential cryptic species we identify are good candidates for further sequencing across the nuclear genome, allowing us to more deeply explore the evolutionary processes in play. Overall, we demonstrate that barcode data, when carefully designed and deployed, are useful both for identifying drivers of divergence and in directing the focus of future genomic studies.

## ACKNOWLEDGMENTS

Thanks for feedback on the manuscript to Courtney Hofman, Hayley Lanier, and JP Masly. We thank the numerous collectors whose specimens were used in both this specific study and the broader Panama Barcoding Initiative, and the institutions at which these vouchers have been deposited. These include the Smithsonian Tropical Research Institute, the Smithsonian Museum of Natural History, the University of Alaska Museum, the University of Washington Burke Museum, the Louisiana State University Museum of Zoology, the Cornell Museum of Vertebrate Zoology, the Field Museum of Natural History, the Academy of Natural Sciences at Drexel University, and the University of Kansas Biodiversity Institute and Natural History Museum.

