## Supplemental Figures S1-S34 for "Comparative phylogeography reveals widespread cryptic diversity driven by ecology in Panamanian birds"

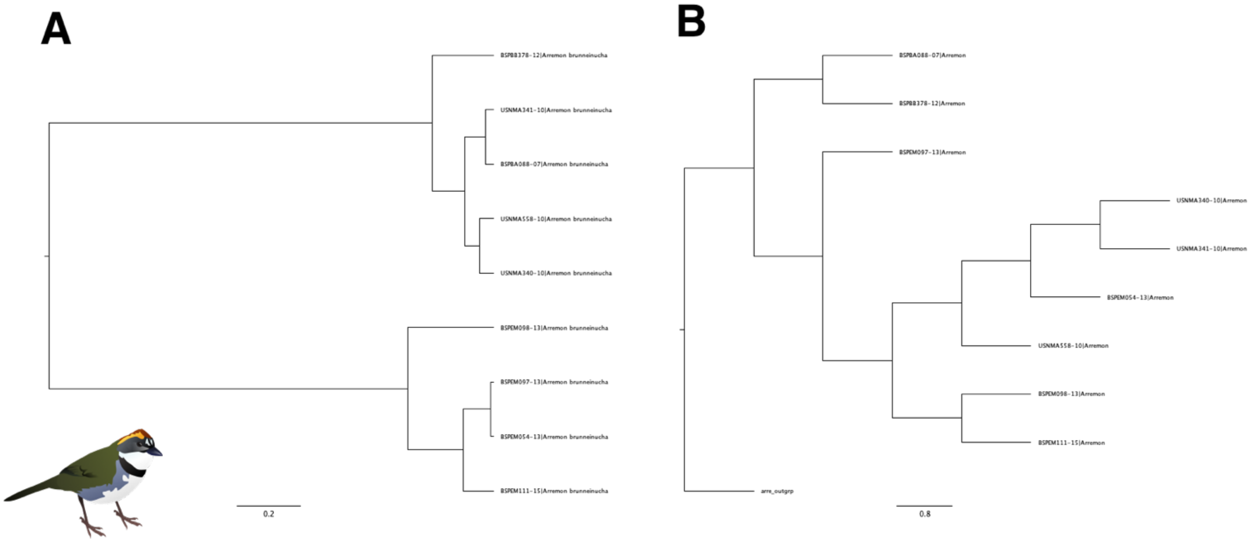


Figure S1: COI trees of *Arremon brunneinucha*, with both A) a NJ tree constructed in MEGA and B) a ML tree constructed in RaxML.

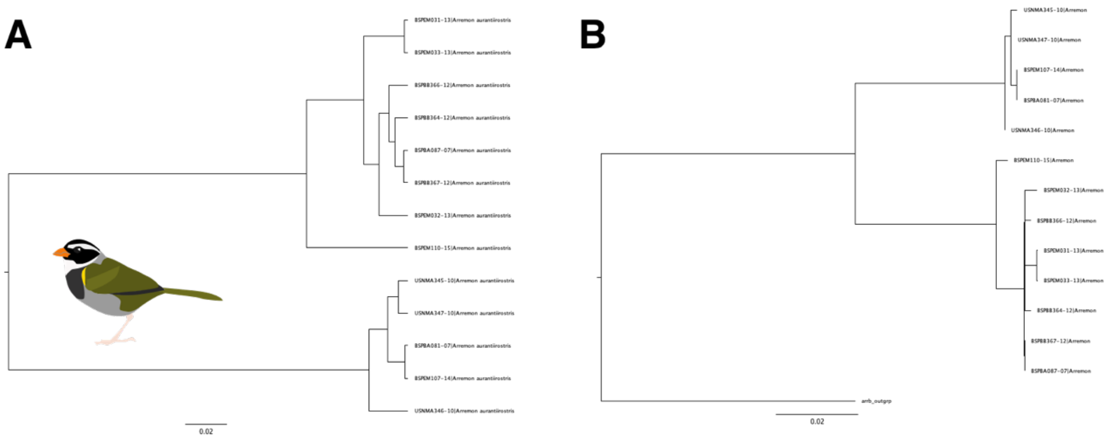


Figure S2: COI trees of *Arremon aurantiirostris* with both A) a NJ tree constructed in MEGA and B) a ML tree constructed in RaxML.

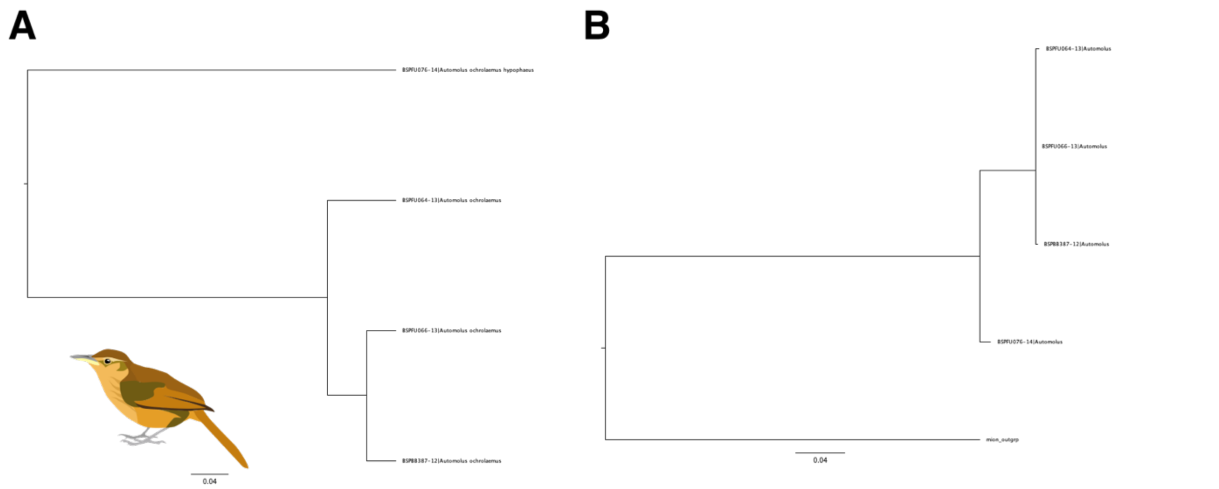


Figure S3: C COI trees of *Automolus ochrolaemus* with both A) a NJ tree constructed in MEGA and B) a ML tree constructed in RaxML.

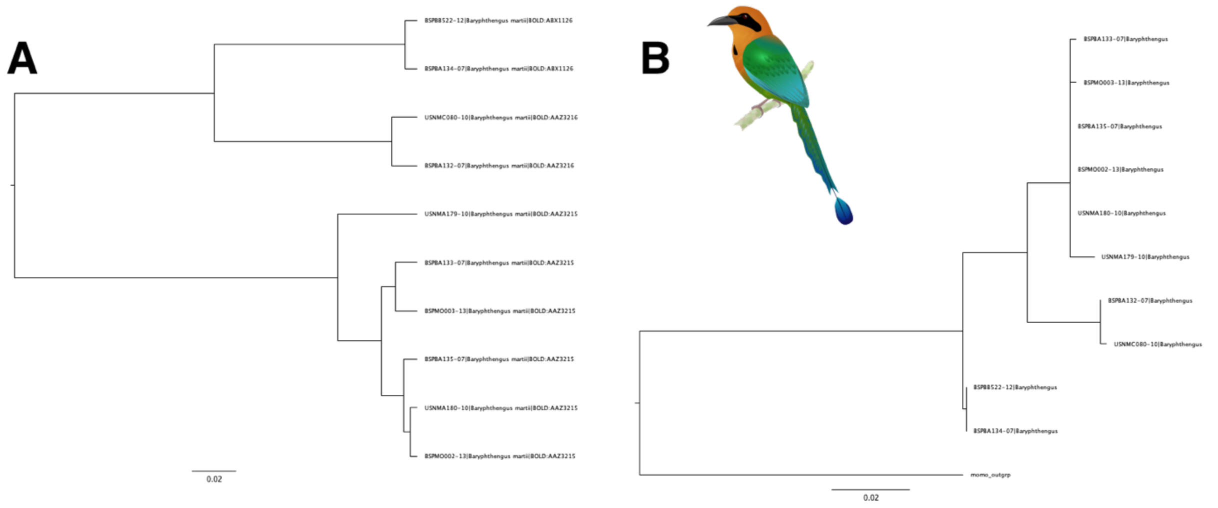


Figure S4: COI trees of *Baryphthengus martii* with both A) a NJ tree constructed in MEGA and B) a ML tree constructed in RaxML.

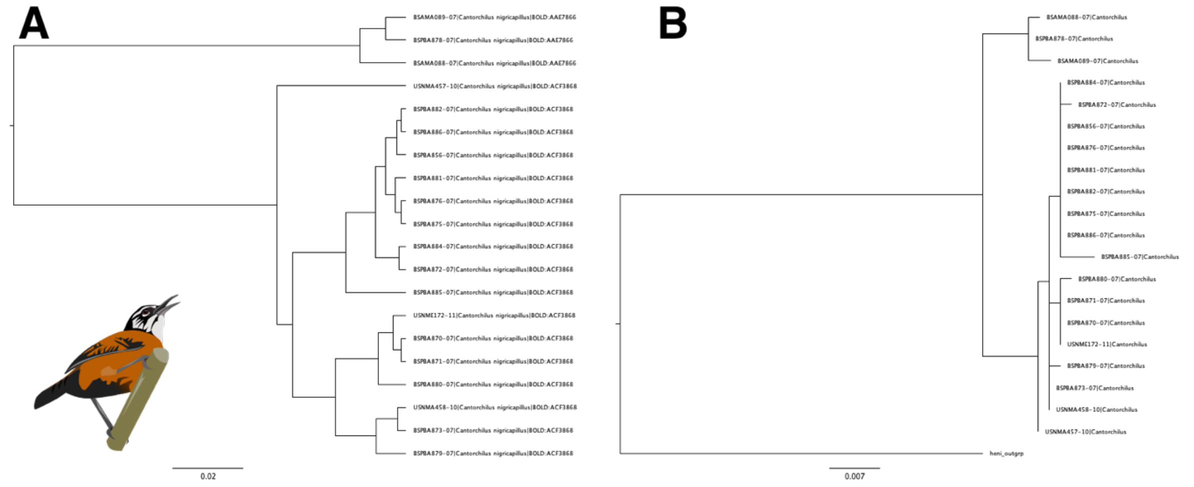


Figure S5: COI trees of *Cantorchilus nigricapillus* with both A) a NJ tree constructed in MEGA and B) a ML tree constructed in RaxML.

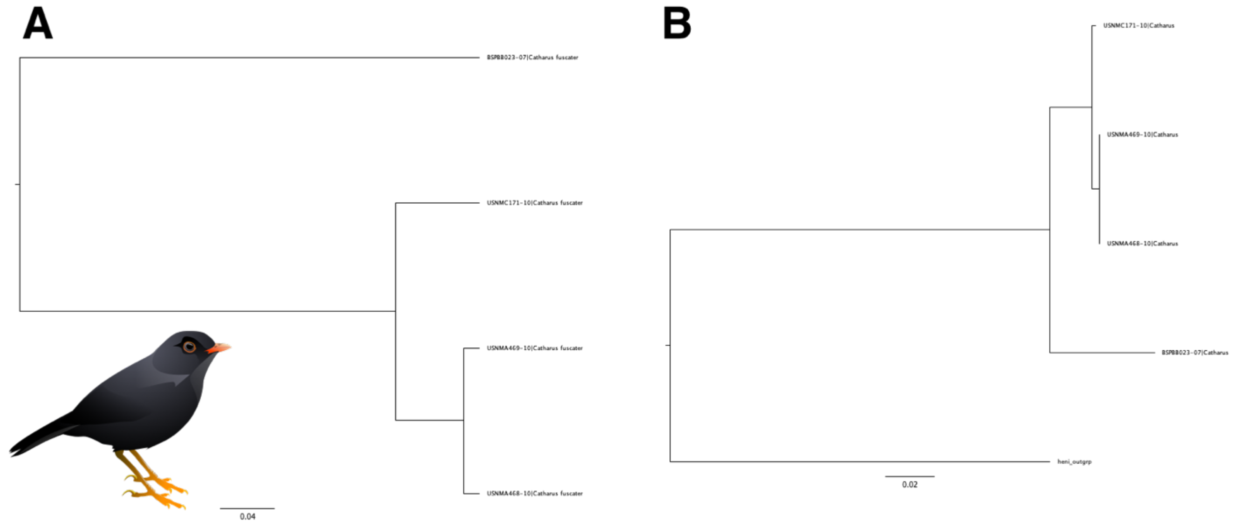


Figure S6: COI trees of *Catharus fuscater* with both A) a NJ tree constructed in MEGA and B) a ML tree constructed in RaxML.

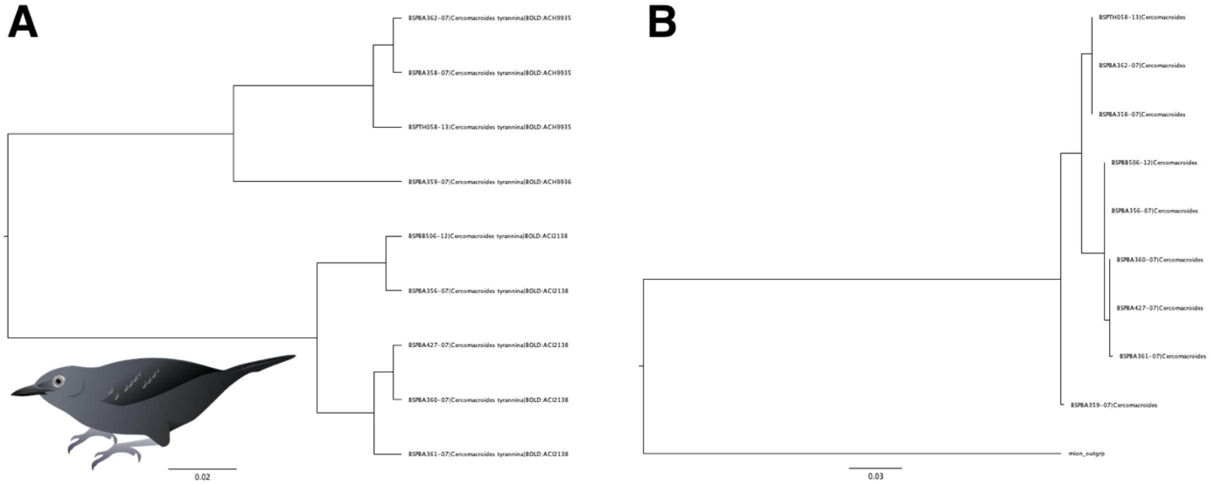


Figure S7: COI trees of *Cercomacra tyrannina* with both A) a NJ tree constructed in MEGA and B) a ML tree constructed in RaxML.

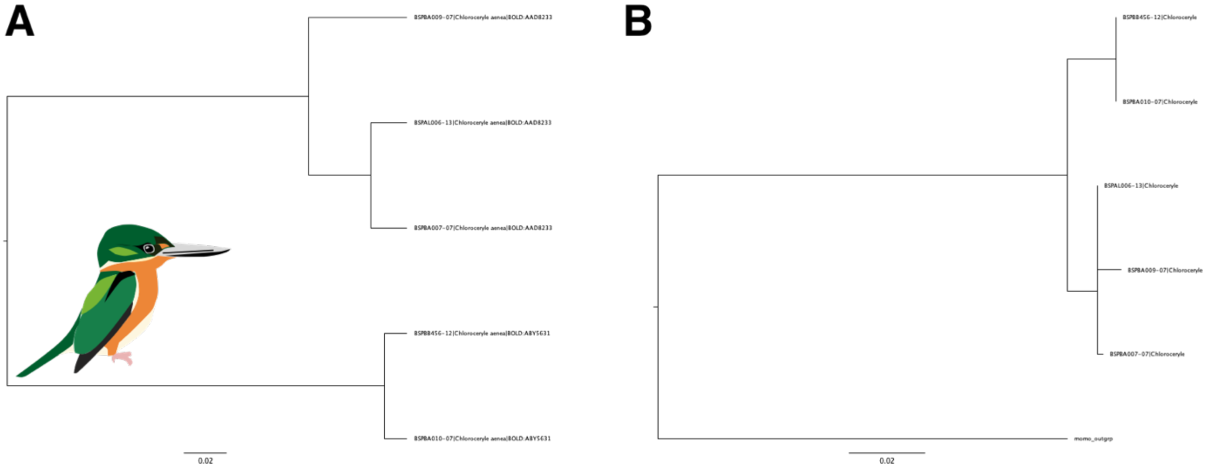


Figure S8: COI trees of *Chloroceryle aenea* with both A) a NJ tree constructed in MEGA and B) a ML tree constructed in RaxML.

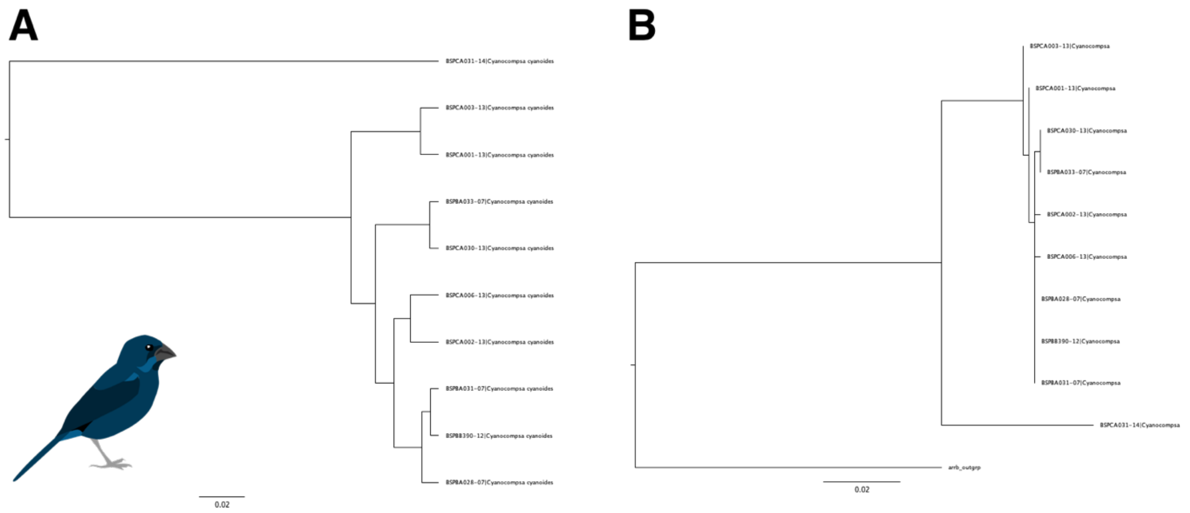


Figure S9: COI trees of *Cyanocompsa cyanoides* with both A) a NJ tree constructed in MEGA and B) a ML tree constructed in RaxML.

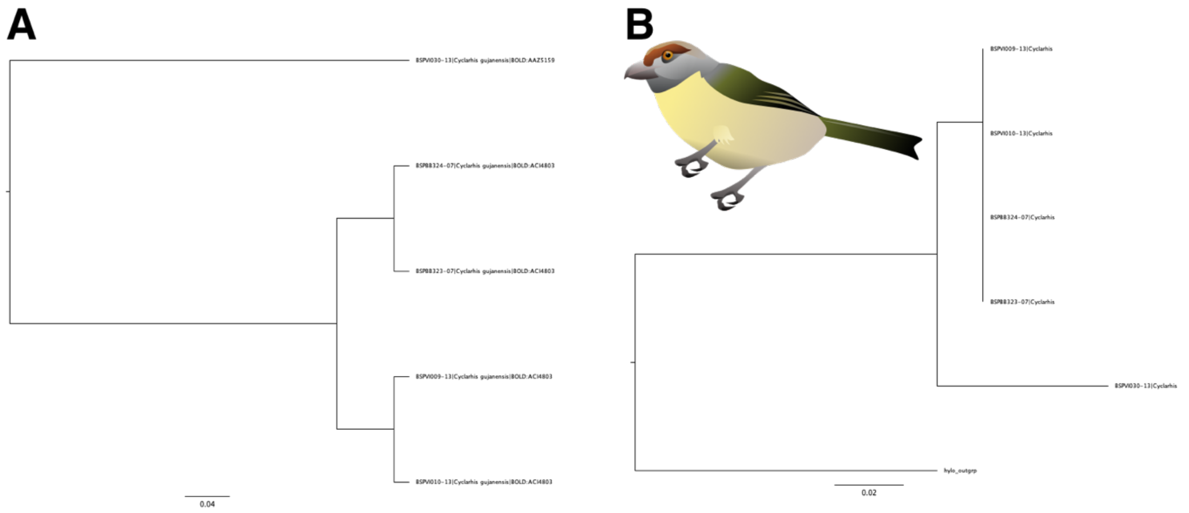


Figure S10: COI trees of *Cyclarhis gujanensis* with both A) a NJ tree constructed in MEGA and B) a ML tree constructed in RaxML.

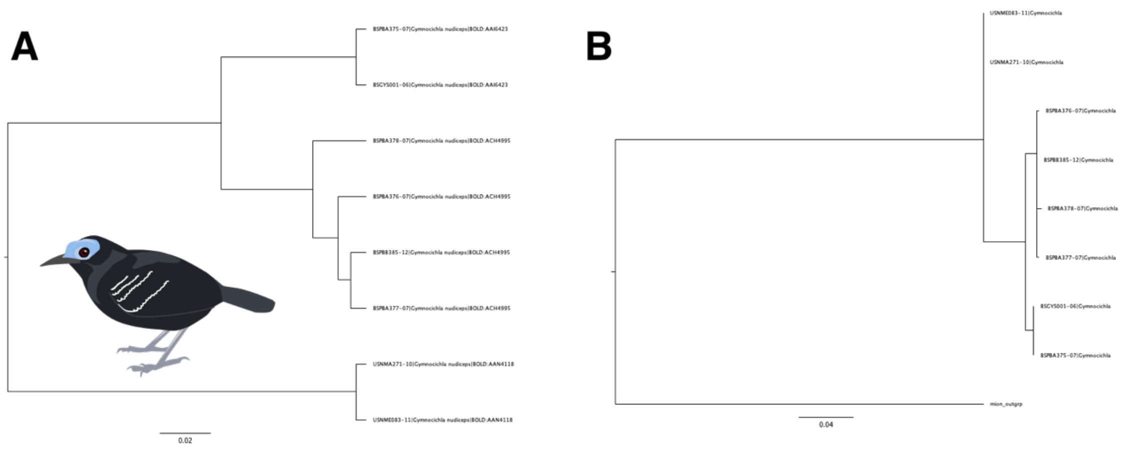


Figure S11: COI trees of *Gymnocichla nudiceps* with both A) a NJ tree constructed in MEGA and B) a ML tree constructed in RaxML.

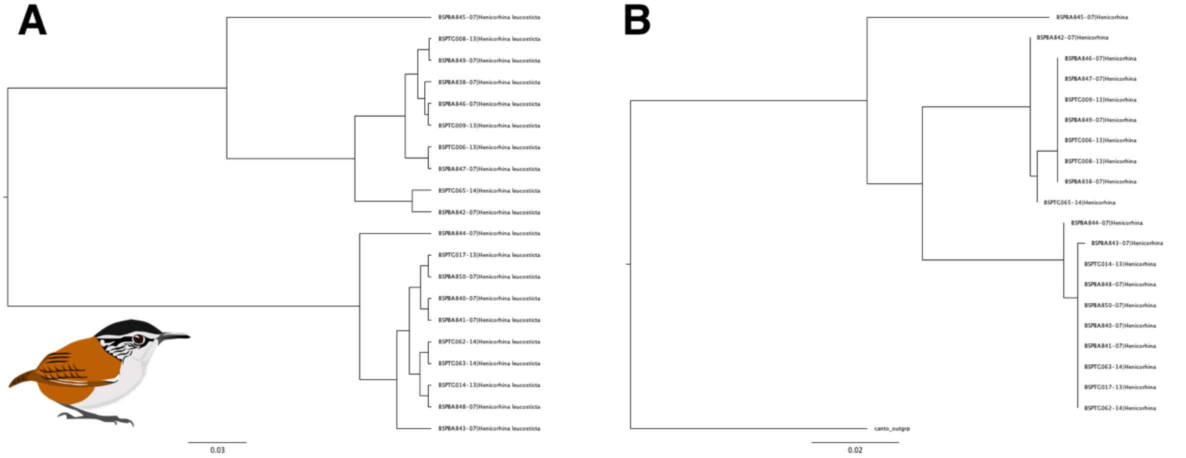


Figure S12: COI trees of *Henicorhina leucosticte* with both A) a NJ tree constructed in MEGA and B) a ML tree constructed in RaxML.

,

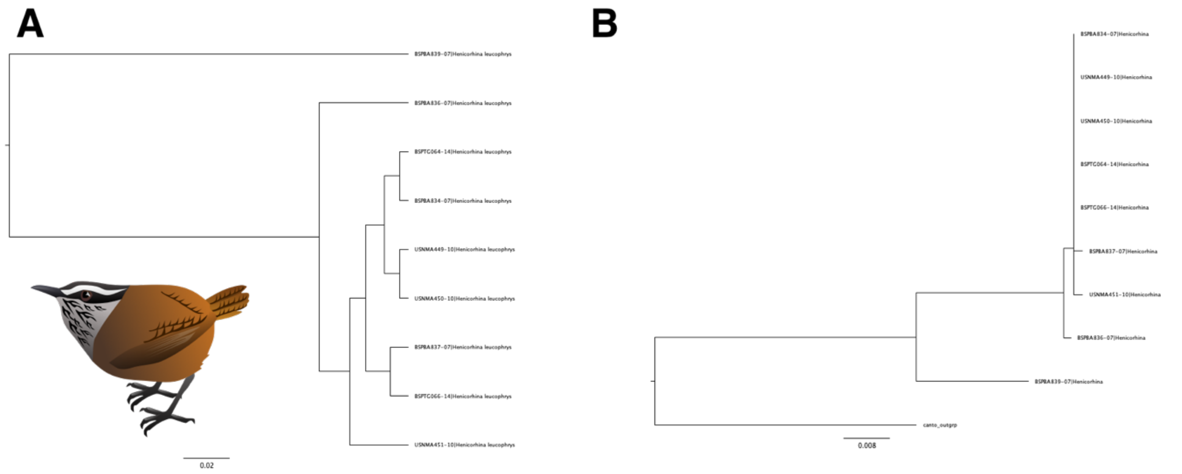


Figure S13: COI trees of *Henicorhina leucophrys* with both A) a NJ tree constructed in MEGA and B) a ML tree constructed in RaxML.

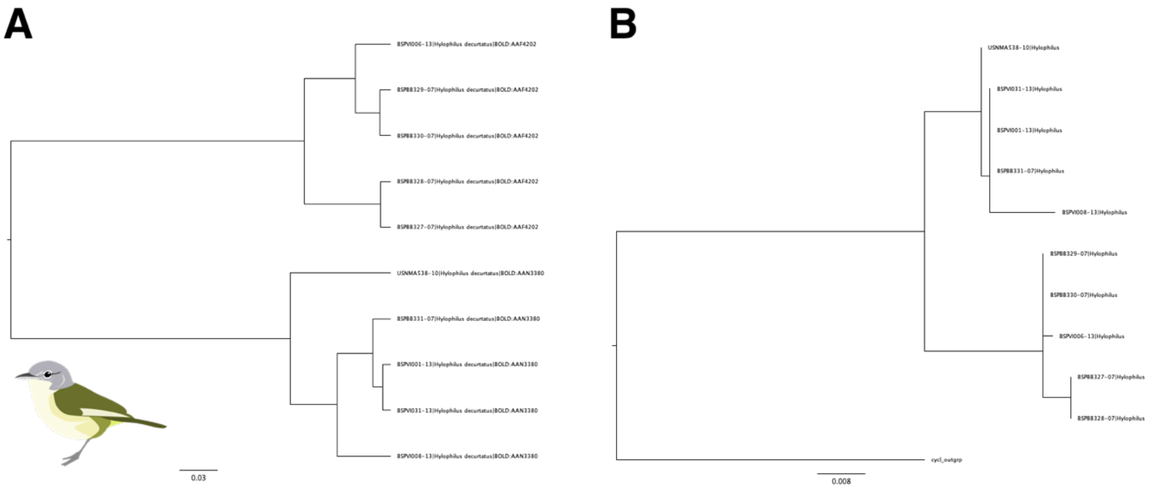


Figure S14: COI trees of *Pachysylvia decurtata* with both A) a NJ tree constructed in MEGA and B) a ML tree constructed in RaxML.

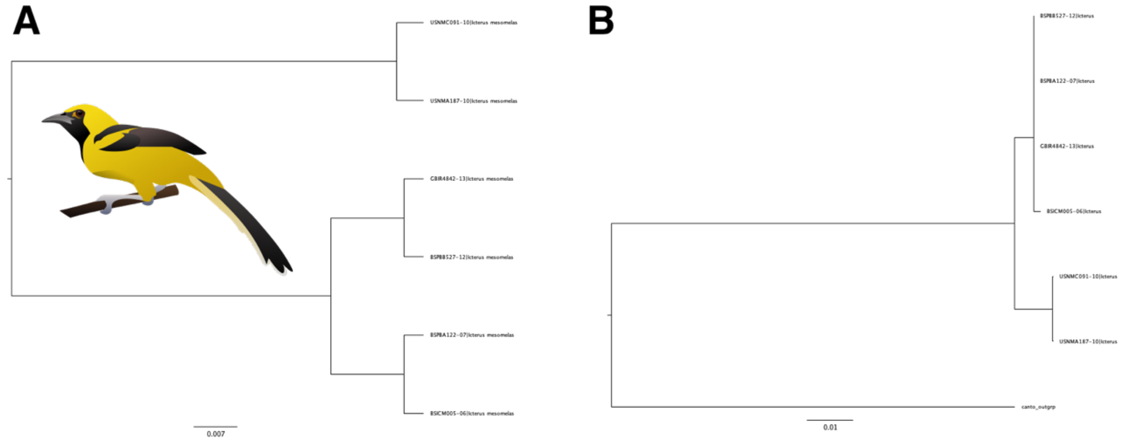


Figure S15: COI trees of *Icterus mesomelas* with both A) a NJ tree constructed in MEGA and B) a ML tree constructed in RaxML.

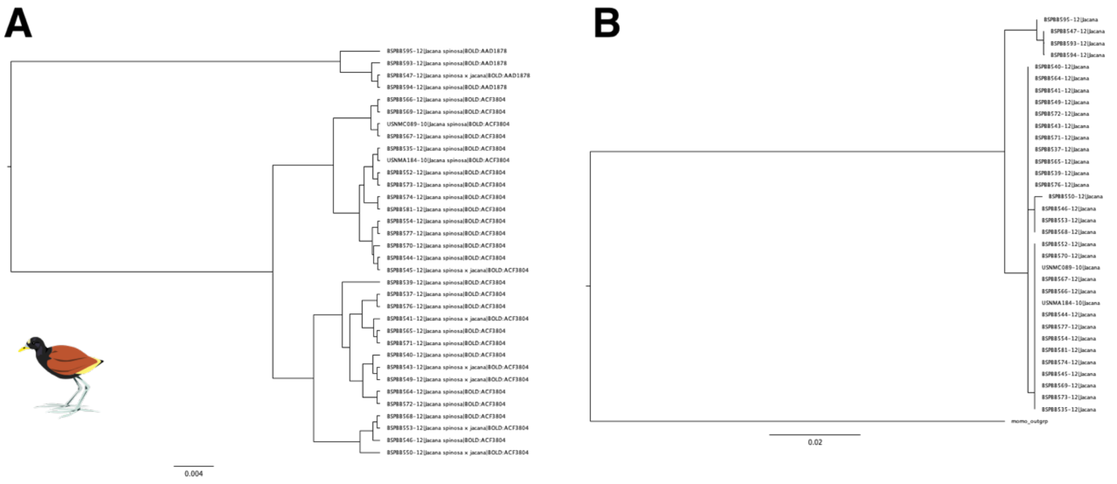


Figure S16: COI trees of *Jacana spinosa* with both A) a NJ tree constructed in MEGA and B) a ML tree constructed in RaxML.

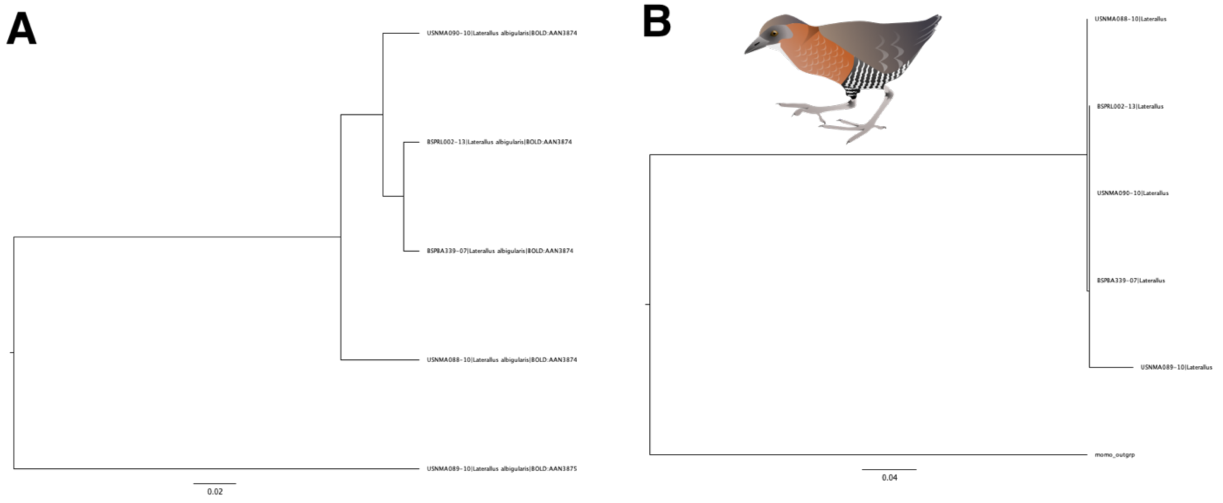


Figure S17: COI trees of *Laterallus albigularis* with both A) a NJ tree constructed in MEGA and B) a ML tree constructed in RaxML.

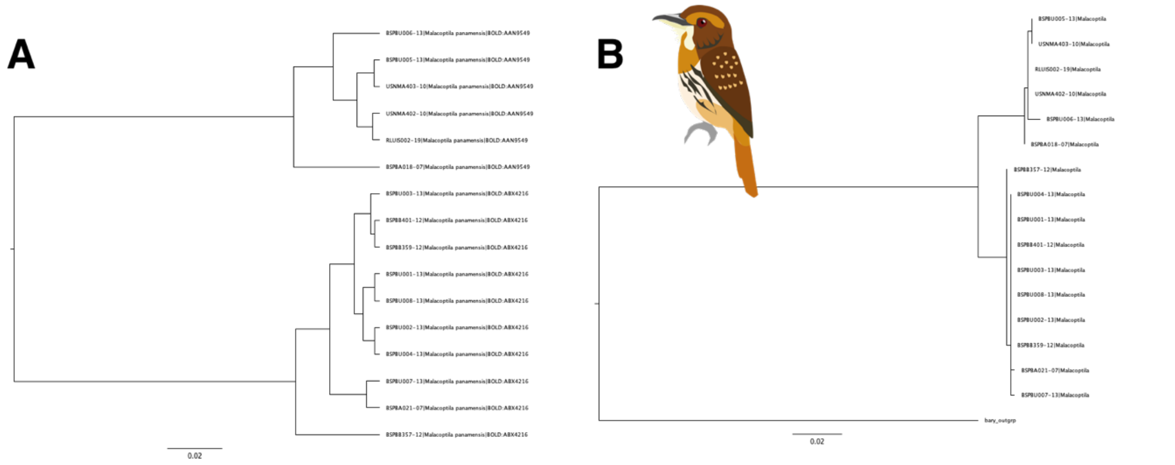


Figure S18: COI trees of *Malacoptila panamensis* with both A) a NJ tree constructed in MEGA and B) a ML tree constructed in RaxML.

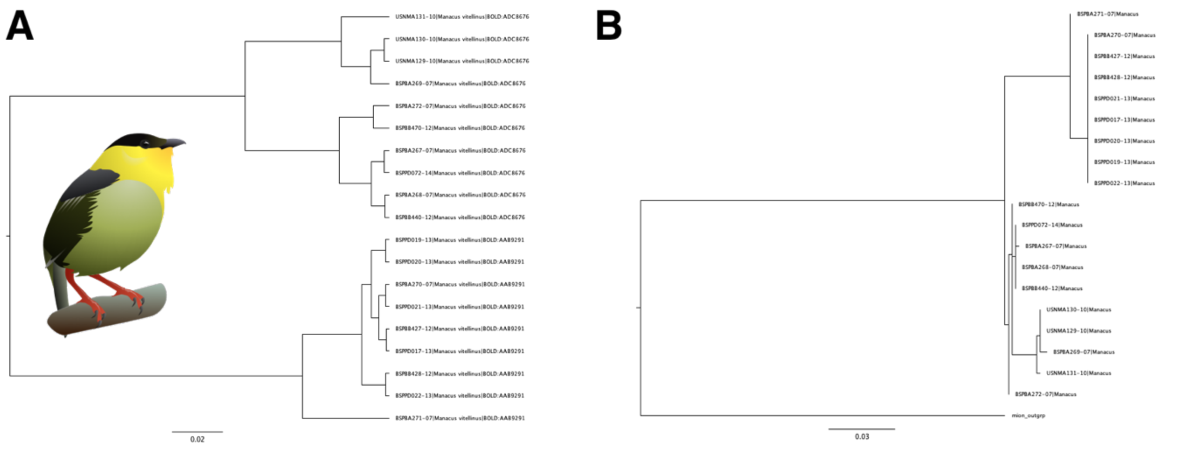


Figure S19: COI trees of *Manacus vitellinus* with both A) a NJ tree constructed in MEGA and B) a ML tree constructed in RaxML.

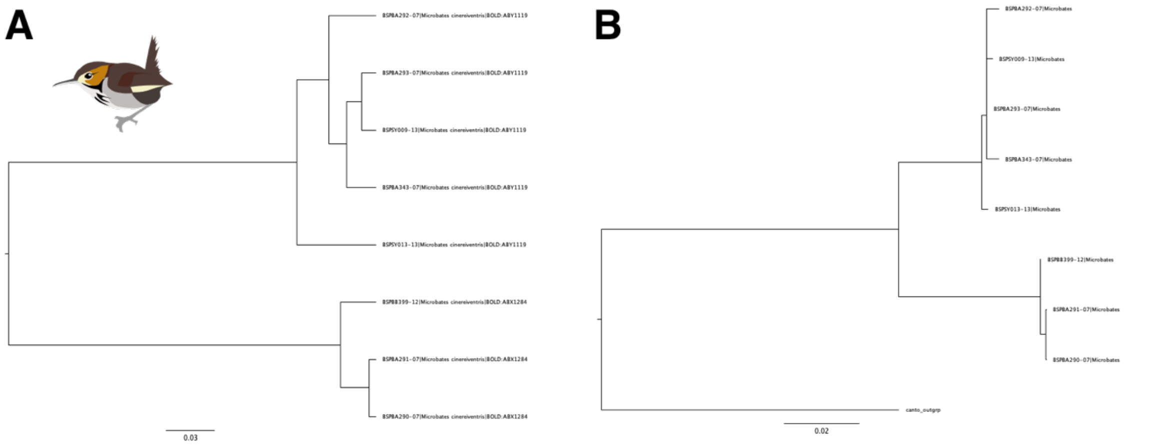


Figure S20: COI trees of *Microbates cinereiventris* with both A) a NJ tree constructed in MEGA and B) a ML tree constructed in RaxML.

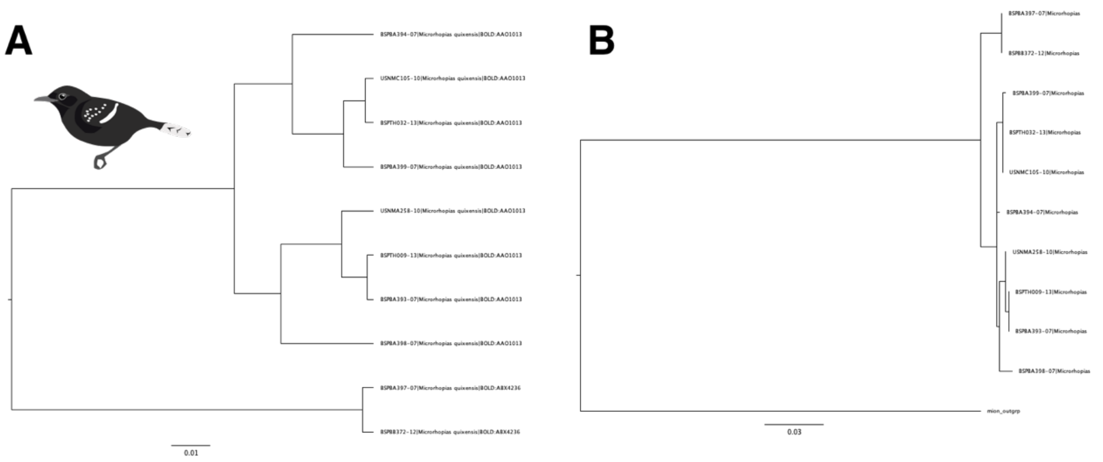


Figure S21: COI trees of *Microrhopias quixensis* with both A) a NJ tree constructed in MEGA and B) a ML tree constructed in RaxML.

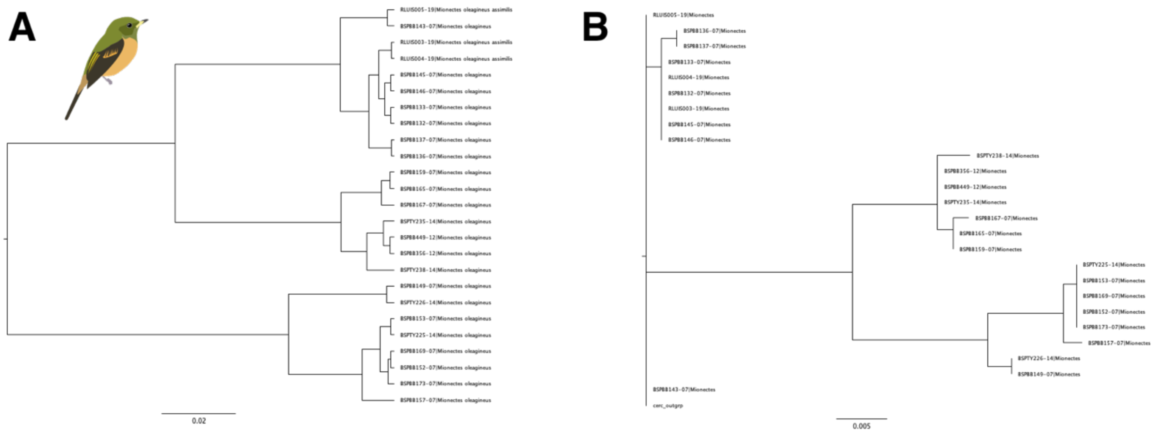


Figure S22: COI trees of *Mionectes oleagineus* with both A) a NJ tree constructed in MEGA and B) a ML tree constructed in RaxML.

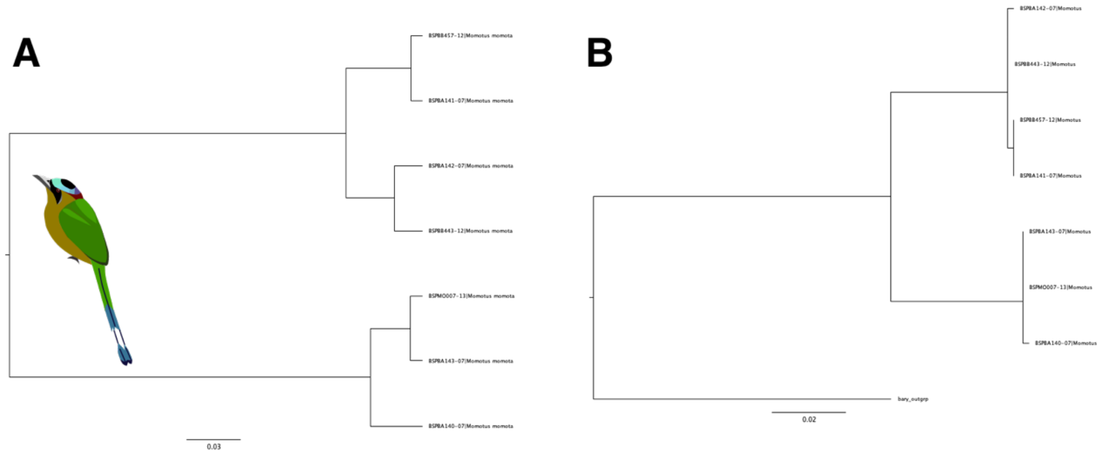


Figure S23: COI trees of *Momotus momota* with both A) a NJ tree constructed in MEGA and B) a ML tree constructed in RaxML.

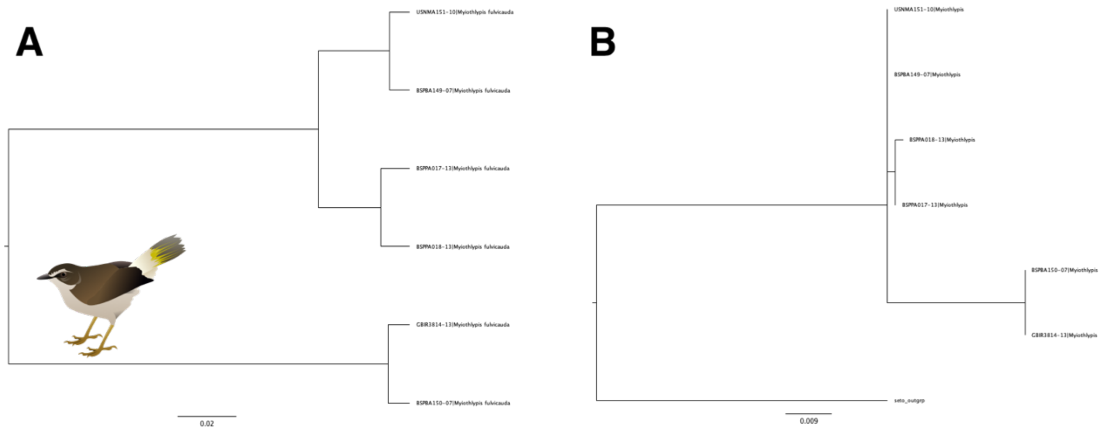


Figure S24: COI trees of *Myiothlypis fulvicauda* with both A) a NJ tree constructed in MEGA and B) a ML tree constructed in RaxML.

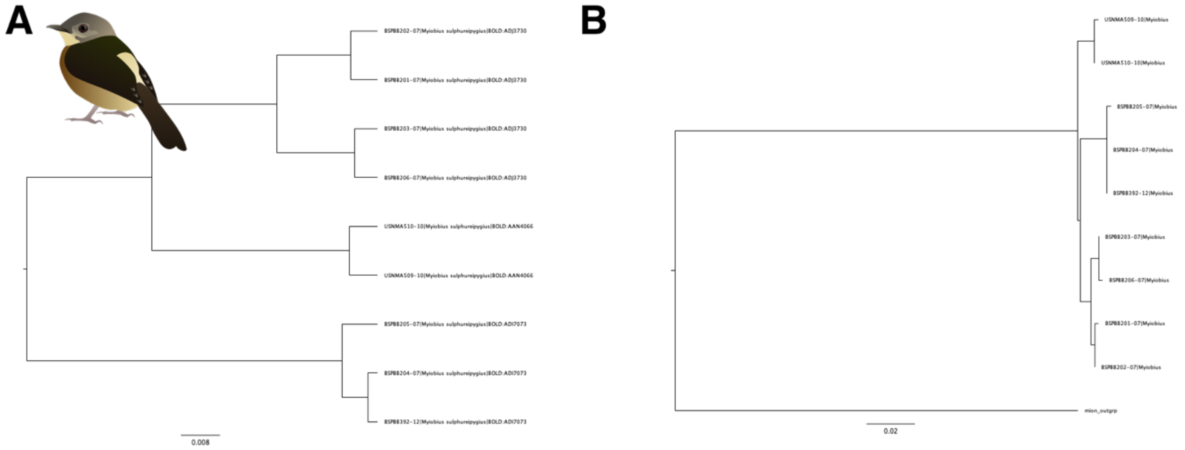


Figure S25: COI trees of *Myiobius sulphureipygius* with both A) a NJ tree constructed in MEGA and B) a ML tree constructed in RaxML.

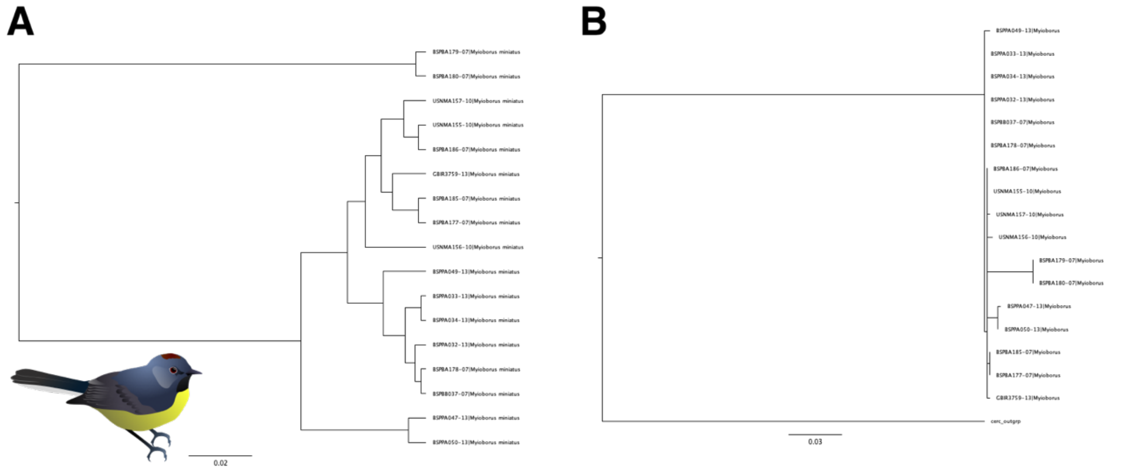


Figure S26: COI trees of *Myioborus miniatus* with both A) a NJ tree constructed in MEGA and B) a ML tree constructed in RaxML.

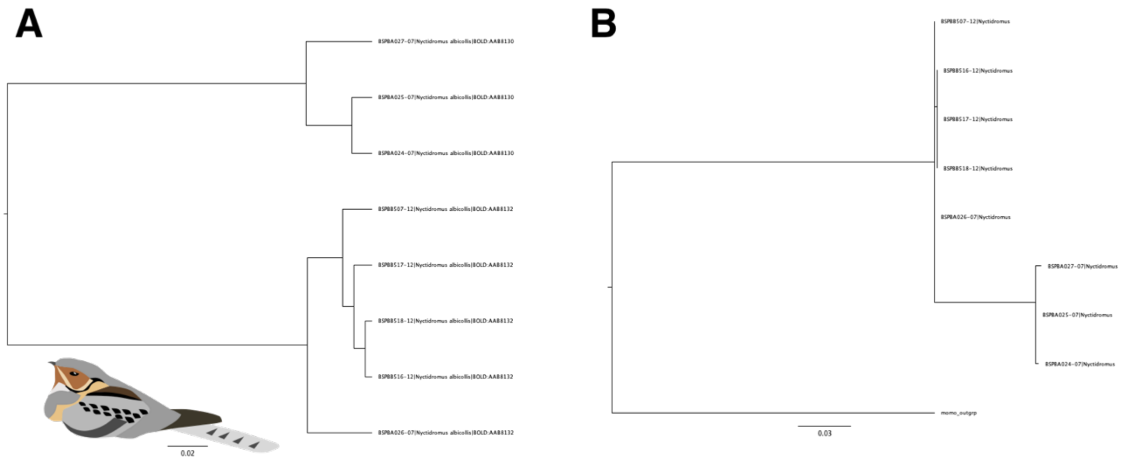


Figure S27: COI trees of *Nyctidromus albicollis* with both A) a NJ tree constructed in MEGA and B) a ML tree constructed in RaxML.

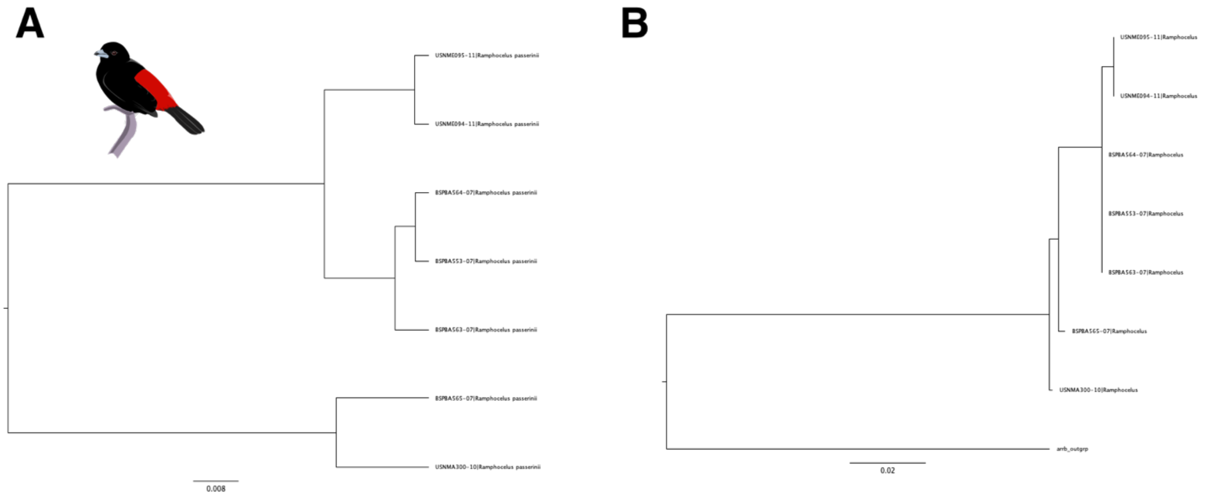


Figure S28: COI trees of *Ramphocelus flammigerus* with both A) a NJ tree constructed in MEGA and B) a ML tree constructed in RaxML.

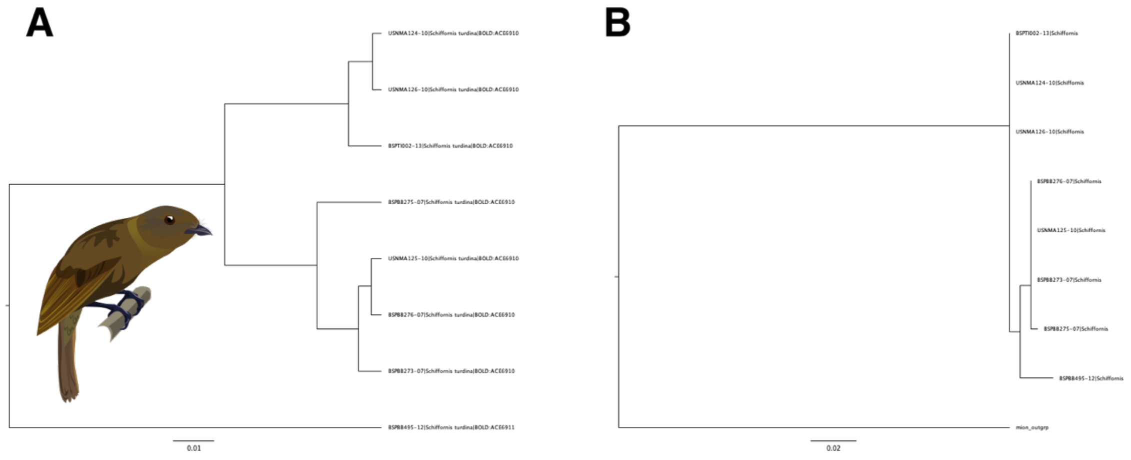


Figure S29: COI trees of *Schiffornis "turdina"* with both A) a NJ tree constructed in MEGA and B) a ML tree constructed in RaxML.

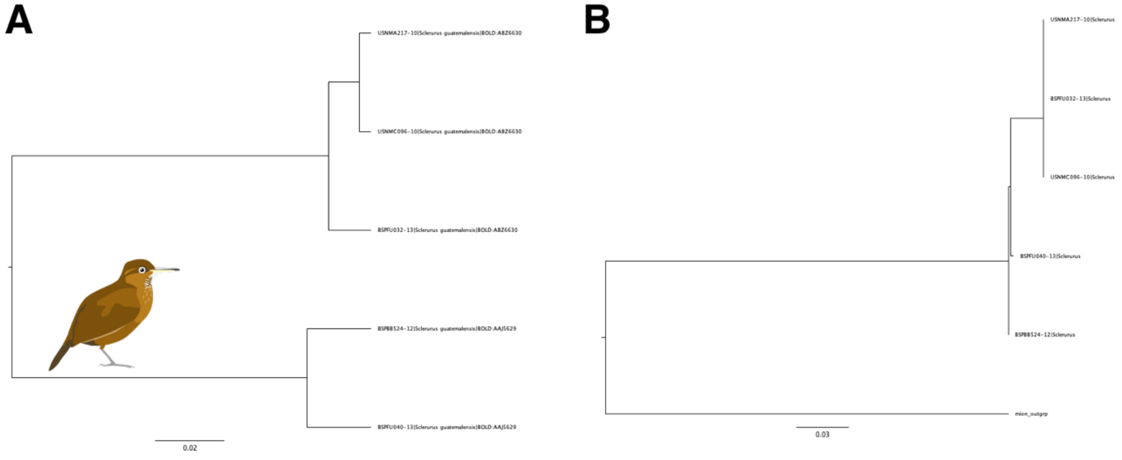


Figure S30: COI trees of *Sclerurus guatemalensis* with both A) a NJ tree constructed in MEGA and B) a ML tree constructed in RaxML.

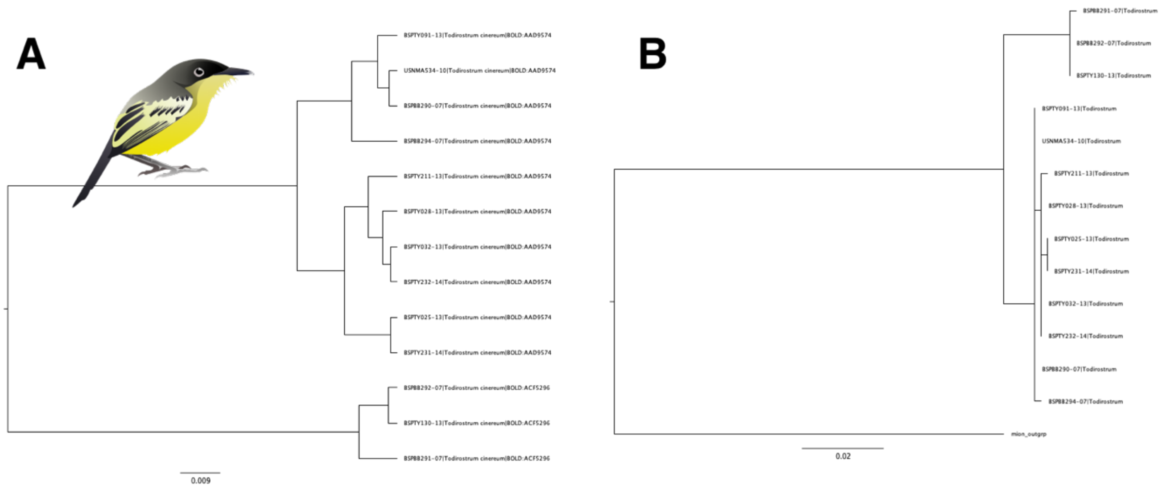


Figure S31: COI trees of *Todirostum cinereum* with both A) a NJ tree constructed in MEGA and B) a ML tree constructed in RaxML.

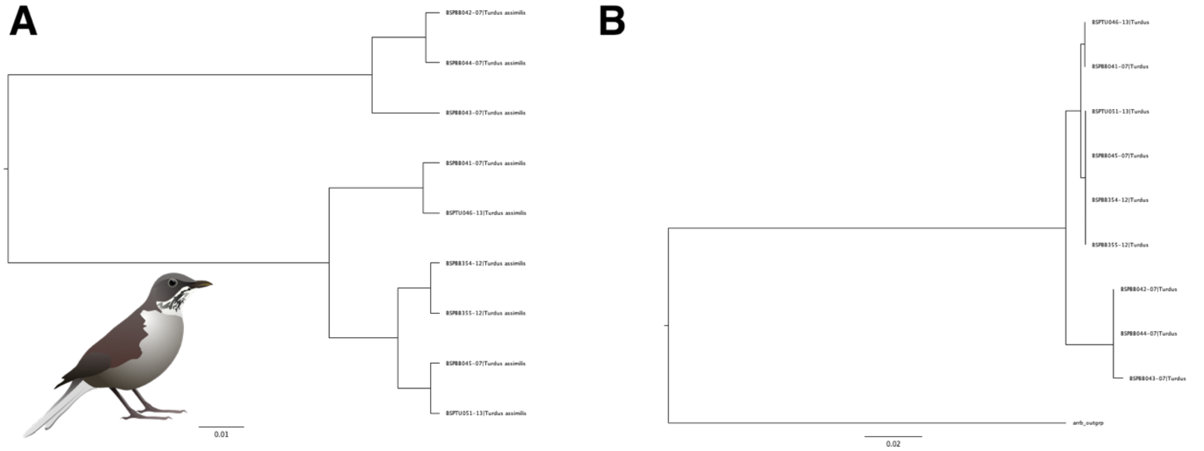


Figure S32: COI trees of *Turdus assimilis* with both A) a NJ tree constructed in MEGA and B) a ML tree constructed in RaxML.

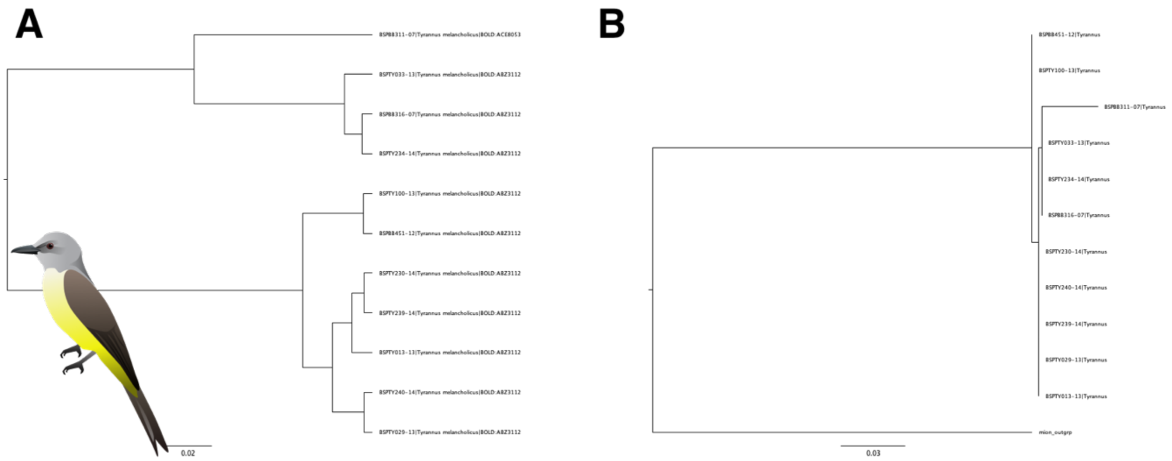


Figure S33: COI trees of *Tyrannus melancholicus* with both A) a NJ tree constructed in MEGA and B) a ML tree constructed in RaxML.

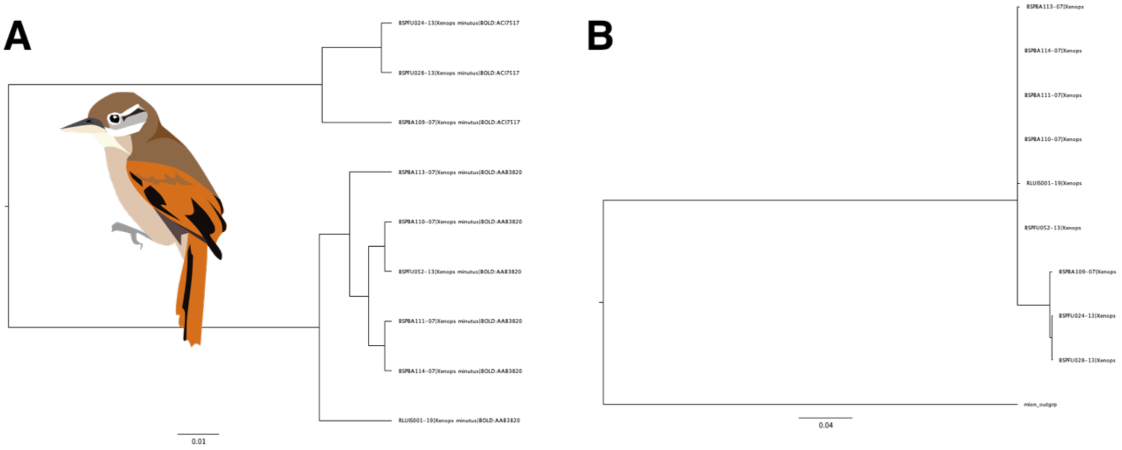


Figure S34 COI trees of *Xenops minutus* with both A) a NJ tree constructed in MEGA and B) a ML tree constructed in RaxML.
